# Identification of Rapaglutin E as An Isoform-specific Inhibitor of Glucose Transporter 1

**DOI:** 10.1101/2025.03.02.641094

**Authors:** Marnie Kotlyar, Zufeng Guo, A.V. Subba Rao, Hanjing Peng, Jingxin Wang, Zhongnan Ma, Cordelia Schiene-Fischer, Gunter Fischer, Jun O. Liu

**Author notes:** Correspondence: Jun O. Liu. Present addresses: Z.G., Basic Medicine Research and Innovation Center for Novel Target and Therapeutic Intervention (Ministry of Education), Department of Breast and Thyroid Surgery of the Second Affiliated Hospital, College of Pharmacy, Chongqing Medical University, Chongqing 400016, China; H.P., US FDA, Silver Spring, MD 20903, United States; J.W., Section of Genetic Medicine, Department of Medicine, Biological Sciences Division, University of Chicago, Chicago, United States.

## Abstract

Natural products rapamycin and FK506 are macrocyclic compounds with therapeutic benefits whose unique scaffold inspired the generation and exploration of the hybrid macrocycle rapafucins. From this library, a potent inhibitor of the facilitative glucose transporter (GLUT), rapaglutin A (RgA) was previously identified. RgA is a pan-GLUT inhibitor of Class I isoforms GLUT1, GLUT3, and GLUT4. Herein, we report the discovery of rapaglutin E (RgE). Unlike RgA, RgE is highly specific for GLUT1. Further characterization revealed that RgE and RgA likely bound to distinct sites on GLUT1 despite their shared FKBP-binding domain, suggesting that the distinct effector domains of RgE and RgA play key roles in recognition of GLUTs.

**TOC GRAPHIC:** 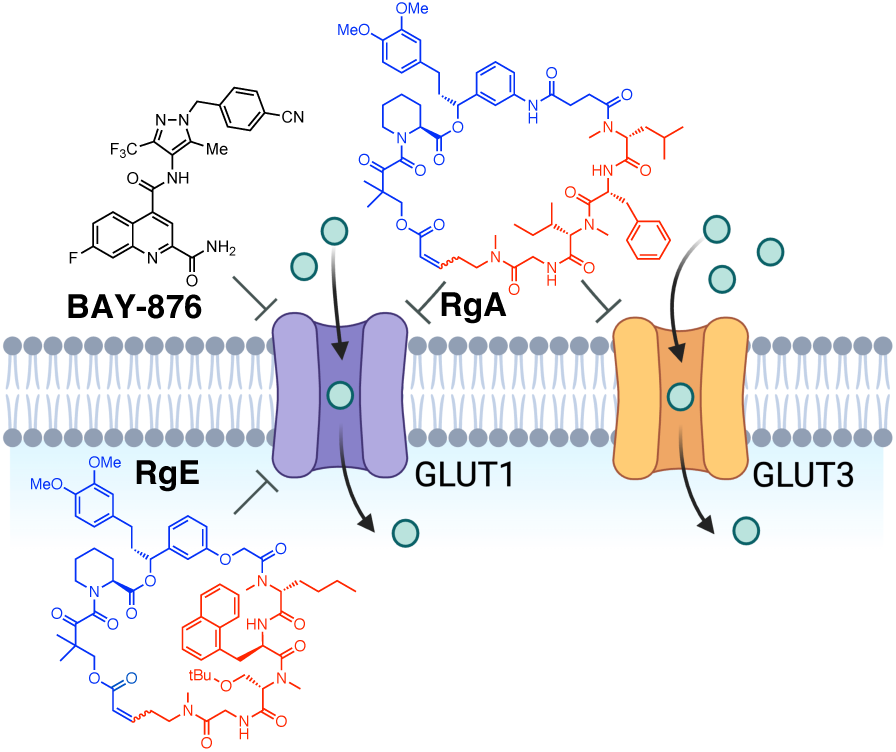

---

Nature serves as a reservoir of bifunctional molecular glues that facilitate protein-protein interactions to modulate biology^1^. Derived from soil bacteria, macrocyclic natural products rapamycin and FK506 are widely used immunosuppressants for organ transplantation and autoimmune diseases^2,3^. Rapamycin also possesses anticancer, antibiotic, anti-ageing and antifungal activity in addition to its immunosuppressive activity. Despite their structural complexity and high molecular weight that are beyond the Lipinski’s rule of 5, both rapamycin and FK506 have been approved for clinical use in their natural forms, suggesting that they offer a privileged drug scaffold. Both rapamycin and FK506 act by recruiting immunophilins FK506-binding proteins (FKBPs) through their shared FK506-binding domains (FKBDs) to form tight binding interactions with their targets, mTOR and calcineurin, respectively^4,5^. The binding to FKBP offers several advantages, including steric hinderance, enhanced stability, intracellular accumulation, and bioavailability^6,7^. Inspired by the unique mode of action of these macrocycles, we designed and synthesized a new FKBD-containing hybrid macrocycle library called rapafucins^8^. Rapafucins contain two versions of an optimized FKBD (FKBD10 or a-FKBD; FKBD11 or e-FKBD) and an oligopeptide-based effector domain^9^. By screening the library in target- and cell-based assays, we have identified novel modulators for difficult-to-drug and undruggable protein targets^8^.

Glucose pays a vital role in biology as it serves as both a key energy source through the production of cellular ATP and a precursor of key building blocks. The high polarity of glucose prevents it from penetrating cellular membranes. It is transported into and out of cells by a family of glucose transporters (GLUT), which shuttles glucose across the plasma membrane through a cycle of conformational changes in a process known as facilitated diffusion^10–14^. There are 14 isoforms of GLUT that span three classes with distinct substrate specificity and tissue distribution. Among them, GLUT1 is the most ubiquitous isoform that is also upregulated in cancer^15,16^. GLUT trafficking and surface expression is dynamic, with extracellular and intracellular signals dictating its production and endosomal transport. Once inside the cell, glucose is broken down via glycolysis into substrates for biosynthesis to support cellular survival and growth. The breakdown of glucose generates pyruvate, which is either converted into lactate or shuttled into the mitochondria as acetyl-CoA where the majority of ATP production and cofactor regeneration occurs through the TCA cycle and oxidative phosphorylation. In 1967, Otto Warburg noted that rapidly proliferating cells such as cancer cells demonstrate an increased demand for glucose to support rapid ATP synthesis via aerobic glycolysis, thus bypassing oxidative phosphorylation in the mitochondria, even though it is a more efficient means of generating ATP per unit of glucose. This upregulation of glycolytic machinery, including GLUT1, and lactate production, independent of oxygen availability, became known as the “Warburg effect”^17,18^. Thus, GLUT serves as an attractive drug target in metabolically dysfunctional cells like cancer for which nutrient limitation can be detrimental^19,20^.

To date a collection of glucose uptake inhibitors have been reported. These comprise multiple classes – natural products, natural product-derived, and synthetic small molecules – with differential potency, selectivity, and mode of action. They include cytochalasin B (cytoB)^21^, WZB-117^22,23^, BAY-876^24^, glutipyran^25^, glutor^26^, and STF-31^19^. CytoB, WZB-117, glutipyran, and glutor are pan-GLUT inhibitors, while BAY-876 and STF-31 are GLUT1-specific. The feasibility of GLUT1 inhibition as cancer treatment has been reported extensively, from [^3^H]-2-deoxy-D-glucose uptake in cancer cells in vitro to tumor xenograft models in vivo^26,27^. However, no GLUT inhibitors have advanced to clinical studies to date.

We previously screened a subset of the rapafucin library for new inhibitors of GLUTs using a 3D rapafucin microarray with cell lysates containing the V5-epitope tagged GLUT1^28^. Rapaglutin A (RgA) was identified as a potent pan-GLUT inhibitor of class I isoforms GLUT1, GLUT3, and GLUT4, with an IC_50_ of 12 nM in a [^3^H]-2-deoxy-D-glucose uptake assay. RgA inhibited glucose uptake in multiple cancer cell lines, leading to a decrease in cellular ATP level, activation of AMP-dependent kinase, and inhibition of mTOR. Moreover, RgA inhibited tumor xenograft growth in vivo without showing toxicity^28^. Aside from the GLUT1-based screen using 3D rapafucin microarray, we have also screened the 3,000-pool, 45,000-compound solution-phase rapafucin library using the alamar blue assay for cell viability in A549 cells (Figure S1). The library was made using partially split-and-pool synthesis with the 15 amino acid building blocks scrambled at the first position of the tetrapeptide effector domain^8^. One of the most potent pools of hits from the FKBD11 or e-FKBD sub-library in the heatmap, E11-72, was further decoded and assessed. Of the 15 rapafucins contained in the pool, only one, E11-72-1 was active (Table S1). E11-72-1 showed dose-dependent inhibition of proliferation of HEK293T cells. Importantly, its activity is affected by glucose level in cell culture medium with greater potency in low-glucose DMEM (Figure S2). This result suggested that E11-72-1 may be competing with glucose, a GLUT substrate, to manifest its anti-proliferative activity. With both E11-72-1 and RgA having an inhibitory effect on A549 proliferation, we then asked whether E11-72-1 affected cell proliferation through its inhibition of GLUT1, like RgA.

Thus, we tested E11-72-1 in a [^3^H]-2-deoxy-D-glucose ([^3^H]-2DG) uptake assay and found that E11-72-1 indeed inhibits GLUT1 with an IC50 of 46 nM (Table S1). Next, an SAR study was undertaken for E11-72-1. We generated analogues bearing amino acid sidechain structural variations within the effector domain (Figure S3, Table S1). E11-72-1-31, in which ^M^Ala in position 4 was replaced with ^M^NIle, showed a modest two-fold improvement in potency. We named this analog Rapaglutin E (RgE) (Figure 1a). RgE exhibited dose-dependent inhibition of [^3^H]-2DG uptake in A549, Jurkat T, PANC10.05, and RBC, with IC50 values in the low-nanomolar range (Figure 1b). Additionally, RgE dose-dependently inhibits proliferation in A549, PANC10.05, HeLa, Jurkat T, and HEK293T cell lines (Figure 1c), demonstrating its broad effect on mammalian cells of different tissue origin.

**Figure 1.**
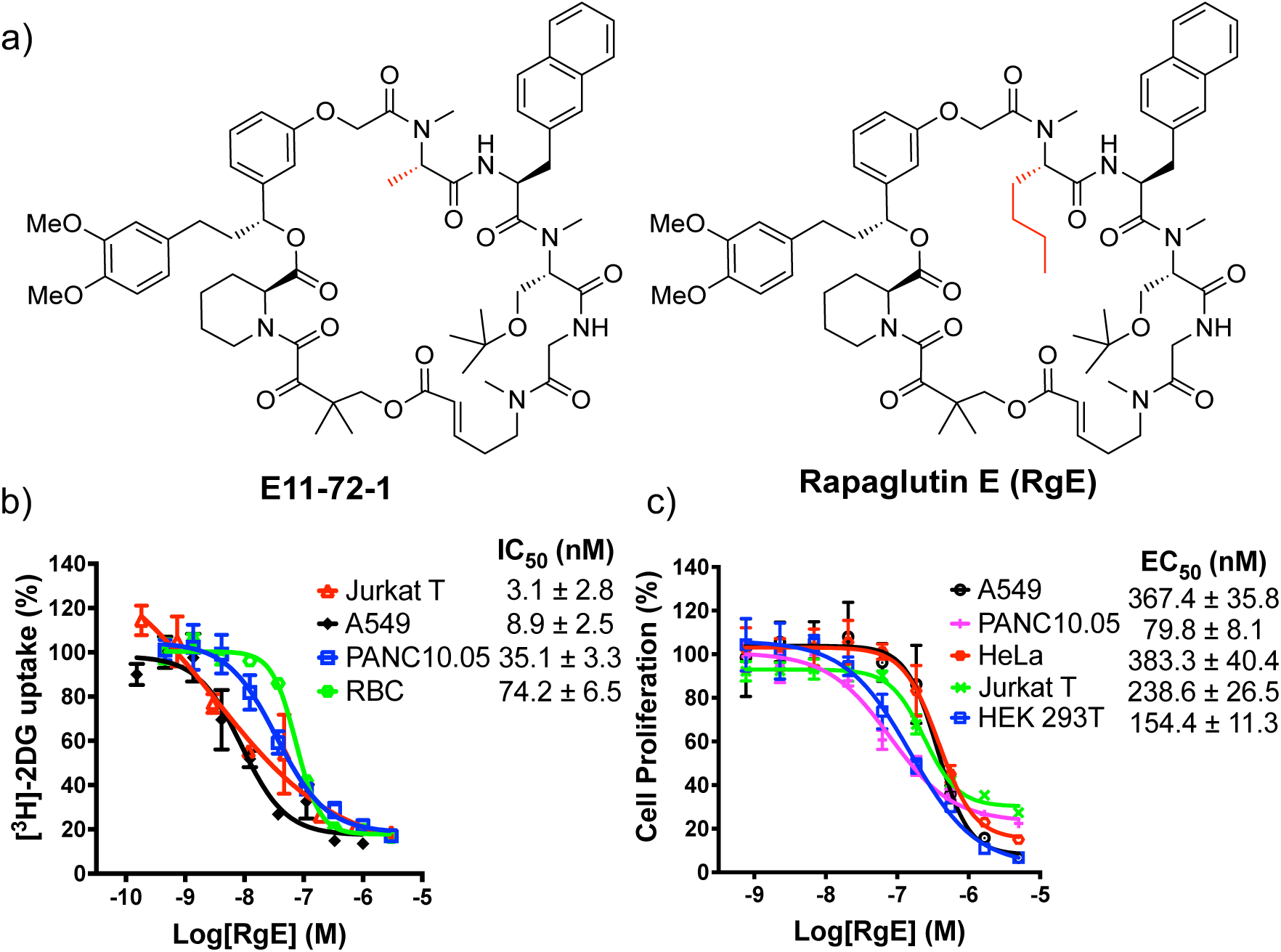
Identification of Rapagutin E (RgE). (a) Chemical structures of E11-72-1 and Rapaglutin E (RgE). (b) Inhibition of 2-deoxy-D-[^3^H] glucose (^3^H-2DG) uptake in A549, PANC10.05, Jurkat T, and red blood cells by RgE. (c) Inhibition of cell proliferation in A549, PANC10.05, HeLa, Jurkat T, and HEK293T cells by RgE (alamar blue assay).

RgE is a 34-membered macrocycle comprised of an e-FKBD, which is two carbons shorter than a-FKBD, bearing an ester rather than an amide linkage between the FKBD and tetrapeptide effector domains. Furthermore, the amino acid composition of RgE’s effector domain is unique and distinct from that of RgA^28^. RgE showed low-nanomolar potency for [^3^H]-2DG uptake inhibition in DLD1 wild type cells (Figure 2a). Given that the effector domain and the ring size of RgE is distinct from RgA, we determined its isoform selectivity using the GLUT1 knockout DLD1 cell line in which GLUT3 is the prevailing isoform for glucose uptake. As expected, RgA that inhibits Class I GLUTs including both GLUT1 and GLUT3 inhibited glucose update in GLUT1 KO cells, albeit with a slightly higher IC50 of 7.8 nM than the WT DLD1 cells (IC50: 4.1 nM) (Figure 2a). In contrast to RgA, the GLUT1 KO cell line became resistant to RgE. This is similar to the known GLUT1-specific inhibitor BAY-876 (Figure 2a). Thus, RgE emerged as a novel, potent GLUT1-specific inhibitor that is structurally and functionally distinct from the pan-GLUT targeting RgA as well as BAY-876. GLUT1 specificity of RgE was further validated using HEK293T WT and HEK293T overexpressing GLUT1, GLUT3, or GLUT4 (Figure 2b). GLUT3 and GLUT4 overexpression conferred resistance to RgE, while GLUT1 overexpressing cells remained sensitive to RgE, albeit with a 7-fold reduction in potency. Together, these results suggest that RgE is a GLUT1-specific inhibitor.

**Figure 2.**
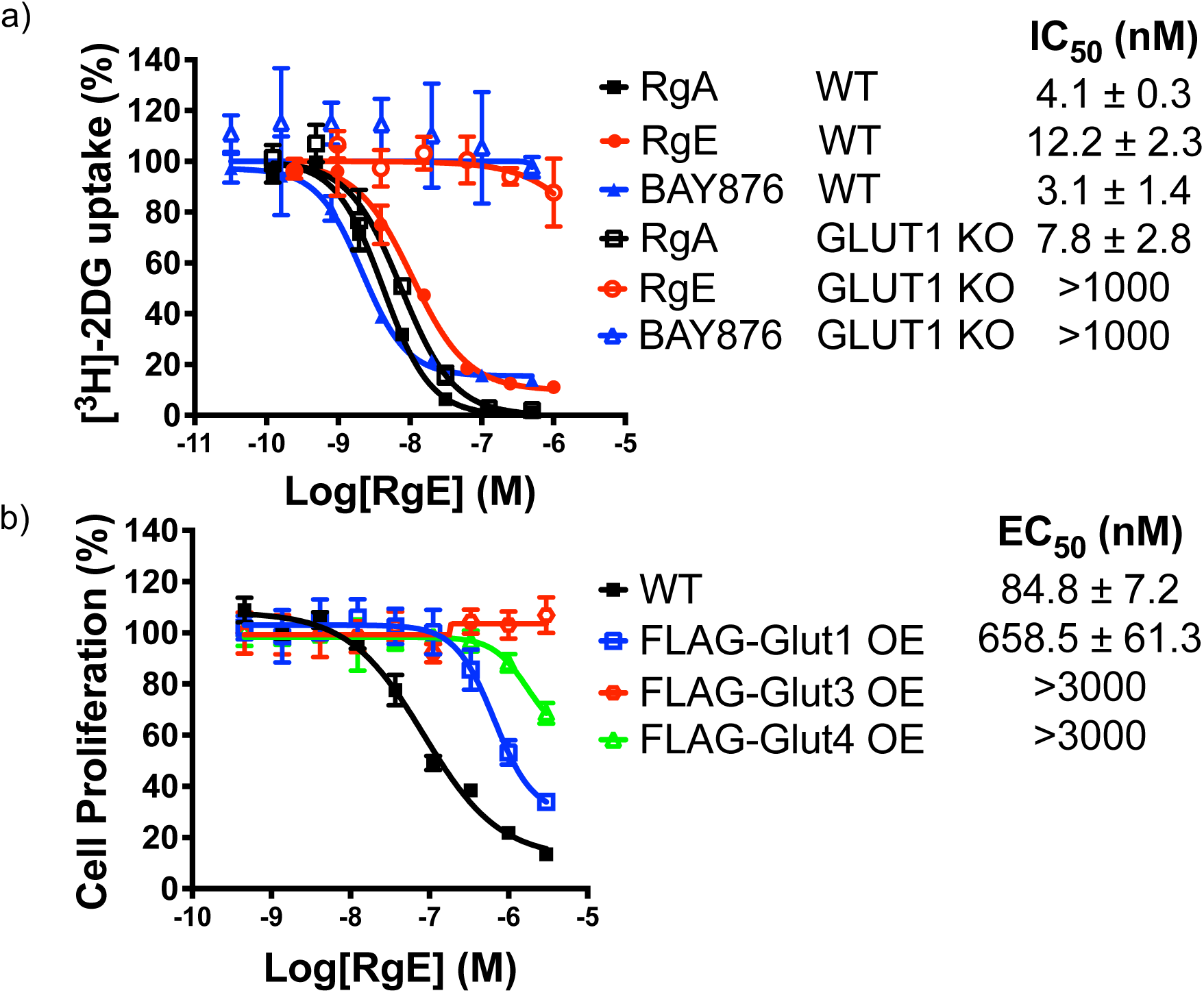
Identification of RgE as a specific inhibitor of GLUT1. (a) Inhibition of [^3^H]-2DG uptake in DLD1 wild type or GLUT1 knock out cells by RgA, RgE, BAY-876, and CytoB; (b) Inhibition of cell proliferation in HEK293T Wild type, GLUT1, GLUT3, or GLUT4 over-expressing cells by RgE (alamar blue assay).

Next, we sought to assess the similarity and differences between RgE and RgA in their interactions with GLUT1. To do so, we decided to apply an affinity pulldown assay using biotin-conjugated probes for RgA^28^ and RgE. For the biotin-RgA conjugate, we have previously attached the biotin moiety to the α,β-unsaturated olefin in the lower linker region of the rapafucin that allowed for retention of activity^28^ (Figure 3a). When we made a similar biotin-RgE conjugate, however, it lost its GLUT1-inhibitory activity. We then attempted an alternative attachment site via the dimethoxyphenyl sidechain (Figure 3a) and the corresponding biotin-RgE conjugate retained significant amount of activity, despite a 40-fold decrease in potency for [^3^H]-2DG uptake inhibition (Figure S4). The difference in sensitivity to biotin attachment at two different sites in the macrocycles suggests that their binding to GLUT1 involves distinct structural elements aside from their unique effector domain, contributing to their different isoform selectivity.

**Figure 3.**
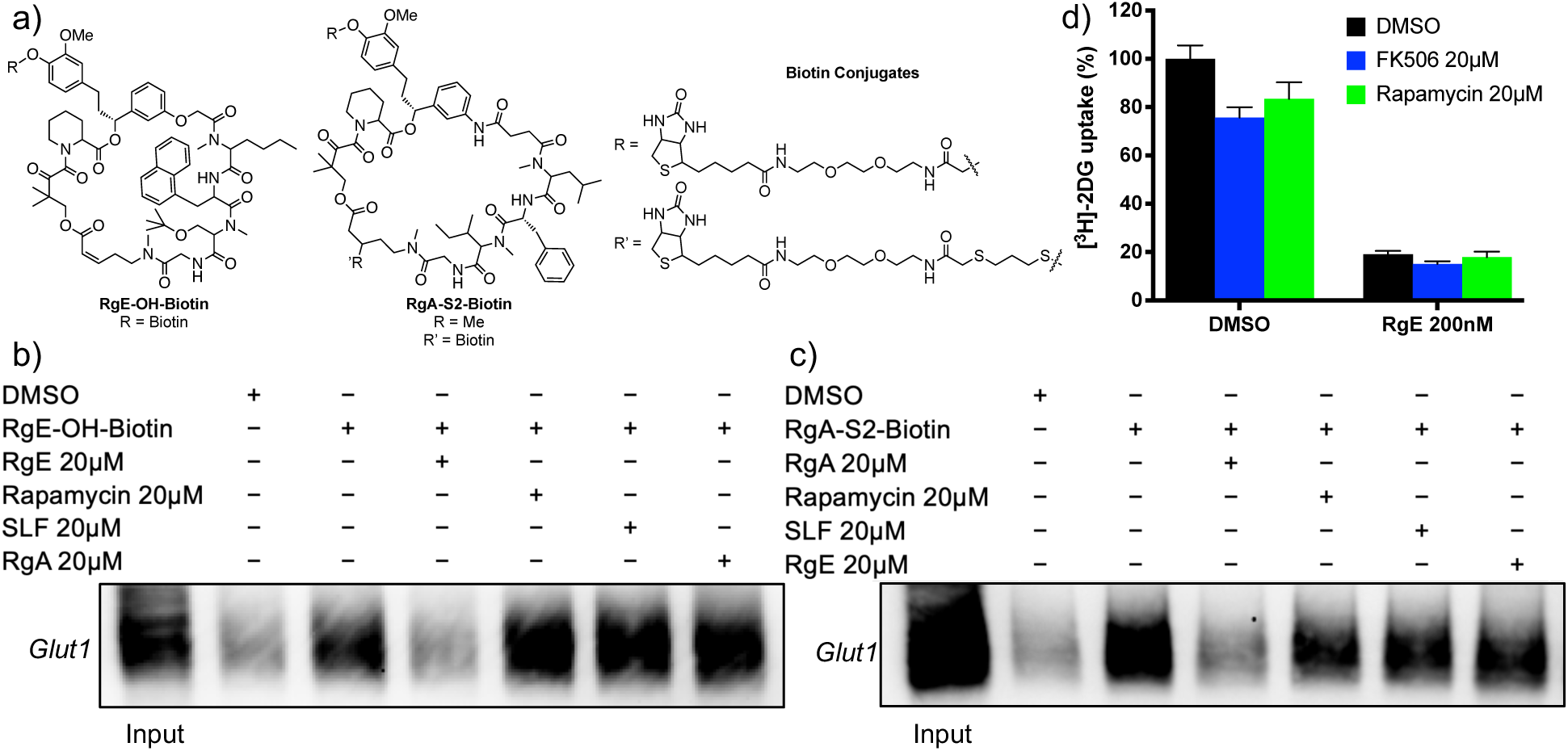
RgE binding to GLUT1 and [^3^H]-2DG uptake inhibitory activity is independent of FKBP. (a) Structures of biotin probes for RgE and RgA. (b, c) Affinity pulldown of detergent-solubilized GLUT1 by a biotin-RgE or biotin-RgA conjugate and competition by free RgE, rapamycin, SLF or RgA. GLUT1 was detected by western blotting; (d) Inhibition of [^3^H]-2DG uptake in A549 cells by 200nM of RgE, 20 μM of FK506, 20 μM of rapamycin and their combinations.

Using the biotin-RgE conjugate, we successfully pulled down detergent-solubilized GLUT1 as judged by Western blot with an anti-GLUT1 antibody (Figure 3b). This pulldown of GLUT1 by biotin-RgE was sensitive to competition by excess RgE, but it is insensitive to competition by RgA. In a reciprocal experiment, we showed that pulldown of GLUT1 with biotin-RgA was sensitive to competition with excessive RgA but not RgE (Figure 3c).

These results suggest that RgE and RgA bind to distinct non-overlapping sites on GLUT1. Given that all rapafucins bind FKBP with high affinity, the question arose as to whether inhibition of GLUT1 by RgE is dependent on FKBP. Neither SLF, a synthetic ligand for FKBP, nor rapamycin affected Biotin-RgE interaction with GLUT1, suggesting that the interaction between RgE and GLUT1 is independent of FKBP. Treatment of RgE in combination with 20 µM FK506 or rapamycin in a [^3^H]-2DG uptake assay did not affect the activity of RgE, confirming that RgE’s effect on GLUT1 functional activity is independent of FKBP (Figure 3d). This is further verified using Jurkat T FKBP12 KO, FKBP51 KO, and FKBP52 KO cells (Figure S5), which showed a minimal change in anti-proliferative activity of RgE in the absence of the FKBPs. Finally, inhibition constants for the peptidyl prolyl cis-trans isomerase activity of different isoforms of FKBP in vitro demonstrate that although RgE binding to FKBP is not essential for its cellular phenotype, RgE is still capable of binding and inhibiting FKBP12, FKBP51, and FKBP52 (Table S2).

To further assess the different modes of interaction between RgE and RgA with GLUT1, we turned to an alternative approach using limited proteolysis. In the 1980s, this method was commonly used to study GLUT1 topology, structure, and potential ligand binding sites before the advent of structural biology techniques like cryo-EM and x-ray crystallography^29–31^. Limited proteolysis at either the exterior face of intact erythrocytes or interior face of alkali-stripped ghosts labeled with [^3^H]-cytoB first revealed differential cleavage patterns and an approximate site for cytoB binding^29^. Comparing differences in fragmentation with exofacial and endofacial ligand labeled transporter indicated that there is a major conformational change when ligand is bound at the external rather than internal site, which results in labeled proteolytic fragments^30^. Thus, we applied both intact HEK293T-GLUT1-FLAG cells and solubilized GLUT1 in lysate to capture target susceptibility to pronase digestion following rapaglutin treatment by SDS-PAGE.

Although derived from the same library, RgE and RgA form distinct contacts with GLUT1, with RgE treatment having little effect on the proteolytic pattern resulting from external pronase digestion with intact cells, like CytoB, suggesting that RgE is an endofacial ligand of GLUT1. Despite their contrasting GLUT1 specificities, RgA and BAY-876 produced analogous proteolytic fragment patterns in experiments using both solubilized GLUT1 and intact cells (Figure 4b), supporting the idea that these ligands bind at the exofacial surface to induce a similar conformational change in GLUT1, thereby exposing pronase-susceptible cleavage sites. Thus, we can conclude that RgA and RgE bind to distinct GLUT1 sites (exofacial and endofacial, respectively) to induce differential conformational changes that affect its susceptibility to proteolytic cleavage.

**Figure 4.**
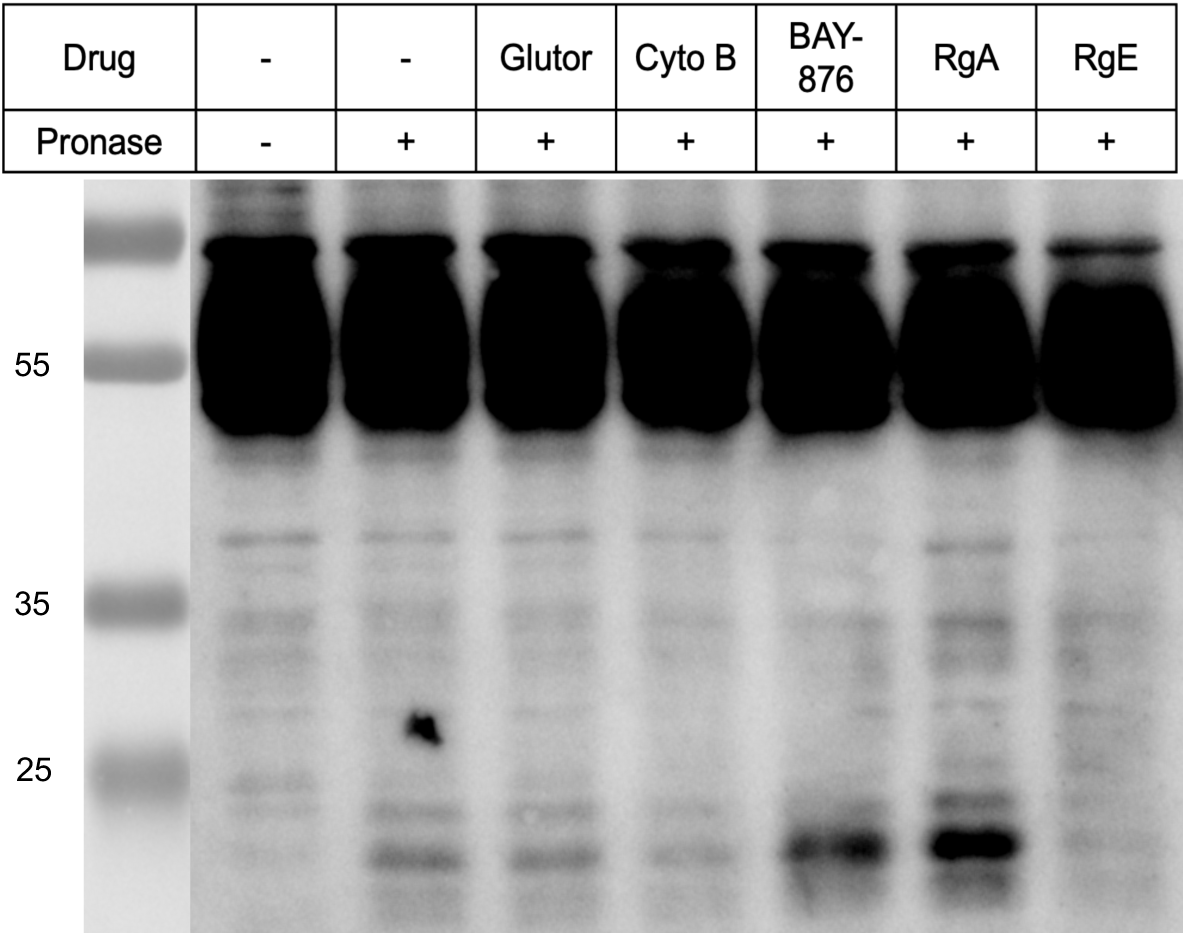
Comparison of GLUT inhibitor effect on GLUT1 limited proteolysis. Western blot of GLUT1 proteolytic fragments produced when HEK293T-GLUT1-FLAG intact cells are pre-treated with 2 µM inhibitor for 30 minutes at 37°C followed by digestion with pronase for 30 minutes at 37°C.

In conclusion, RgE is a novel and highly potent GLUT1-specific inhibitor. Although it shares a common core FKBD structure with RgA, it showed distinct mode of interaction with GLUT1 as reflected in its SAR and the differential protease digestion patterns in comparison with RgA. Together, RgE and RgA expand our arsenal of chemical tools for probing the structure, function and chemical modulation of GLUTs. Unlike other known GLUT inhibitors, RgE and RgA may benefit from favorable PK/PD in vivo due to their high-affinity for and reversible binding to FKBP^7^. Moreover, combination of GLUT1 inhibitors with anticancer drugs or checkpoint inhibitors have shown promise as a therapeutic strategy^32,33^, thus offering exciting potential applications for RgE and RgA.

## Supporting Information

Materials, methods, and characterization data including NMR and mass spectra.

## ACKNOWLEDGMENTS

This work was supported by NIGMS (R01 GM137319), FAMRI (to J.O.L.) and the Cancer Center Support Grant (NCI P30CA006973). M.K. was supported in part by the Chemistry Biology Interface graduate training grant (T32 GM080189). H.P. was a Damon Runyon Fellow supported by the Damon Runyon Cancer Research Foundation (DRG-2191). Special thanks to Dr. Safiat Ayinde for technical guidance and advice.

## SUPPLEMENTARY FIGURES

**Figure S1.**
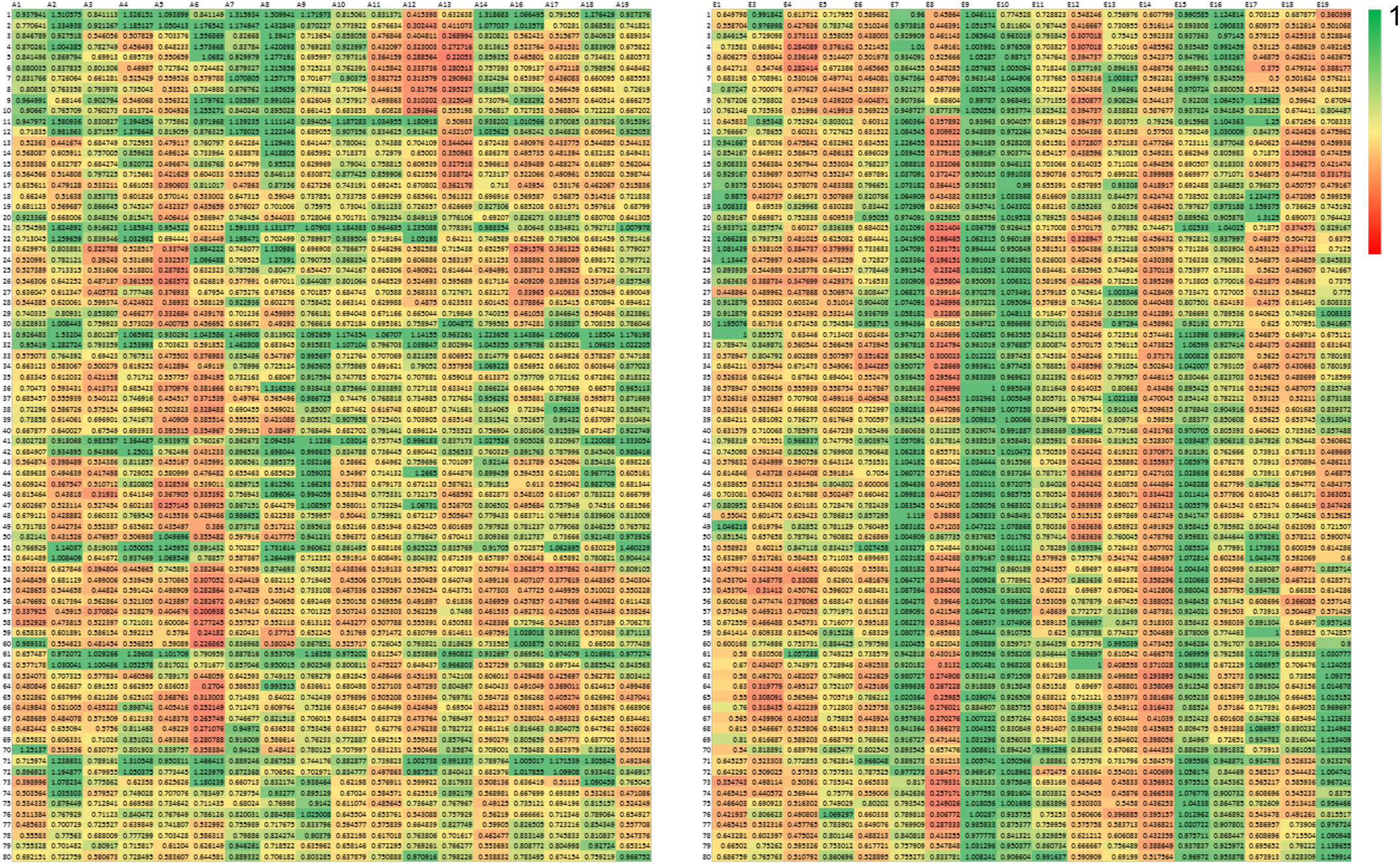
Heatmap of the outcome of the screening of rapafucin libraries using alamar blue assay in A549 cells. FKBD10 or a-FKBD-(left) and FKBD11 or e-FKBD-containing libraries (right) were screened at a final concentration of 3μM. Scale: 1 green, no inhibition; 0 red, 100% inhibition. The arrow points to one of the most potent pool of hits at Row 72, Column E11, that was further decoded.

**Figure S2.**
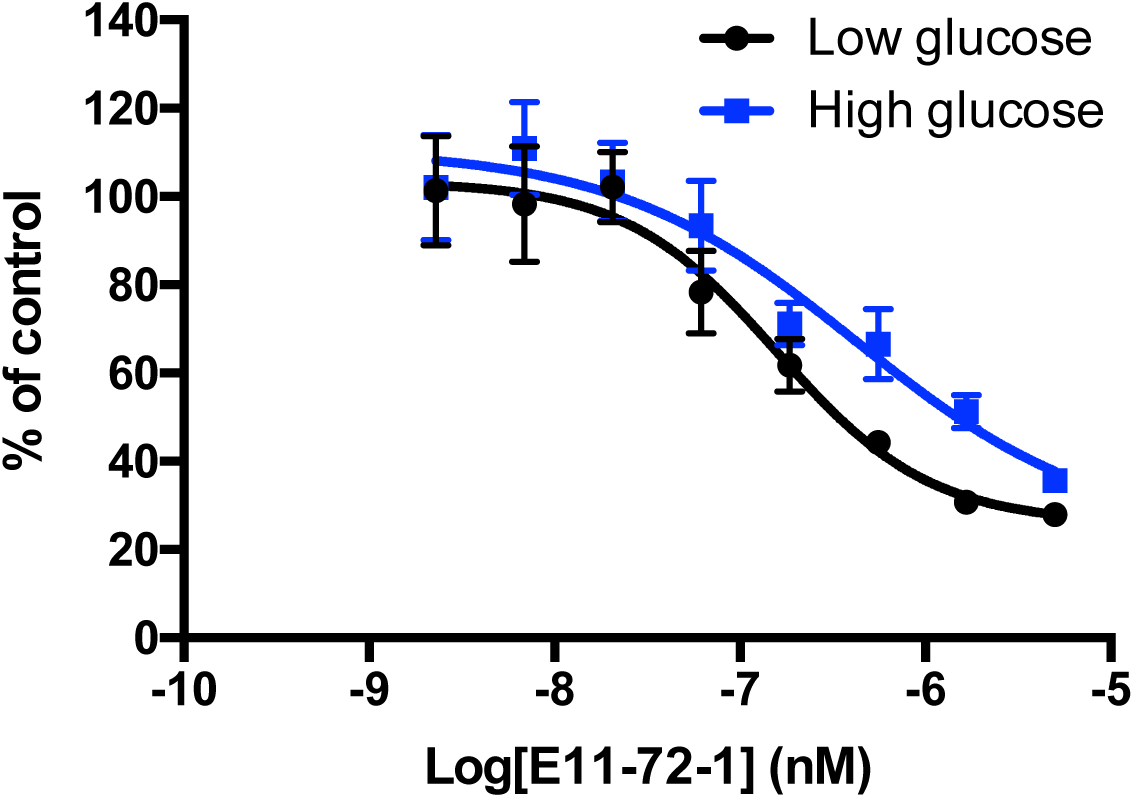
Inhibition of cell proliferation in HEK293T cells by RgE (**8**) in DMEM medium with different glucose concentrations (alamar blue assay). Error bars represent s.d.; data are mean ± s.d.

**Figure S3.**
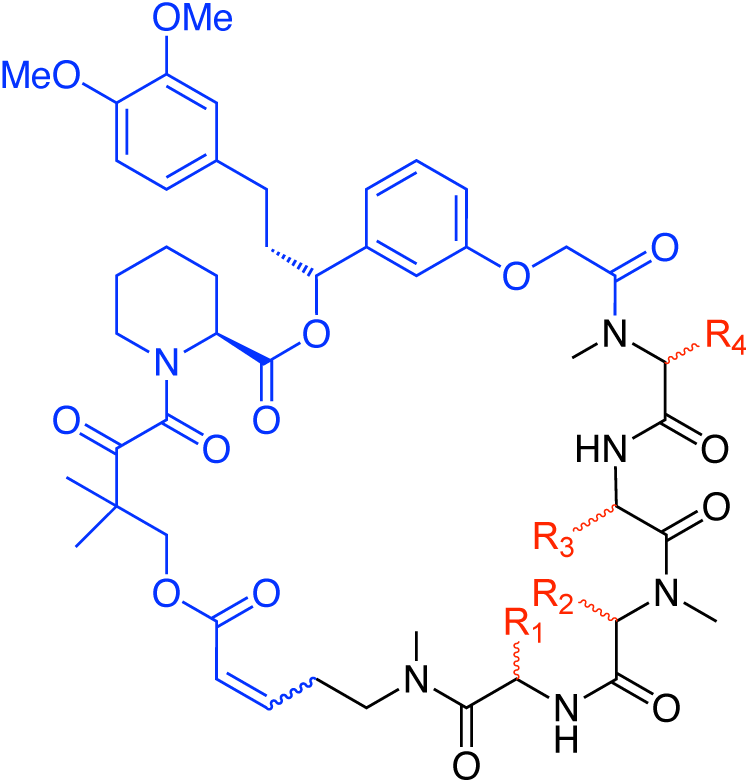
Structure of generic rapafucin with FKBD (**5**) or e-FKBD backbone and an effector domain with variation in its amino acid sidechains.

**Figure S4.**
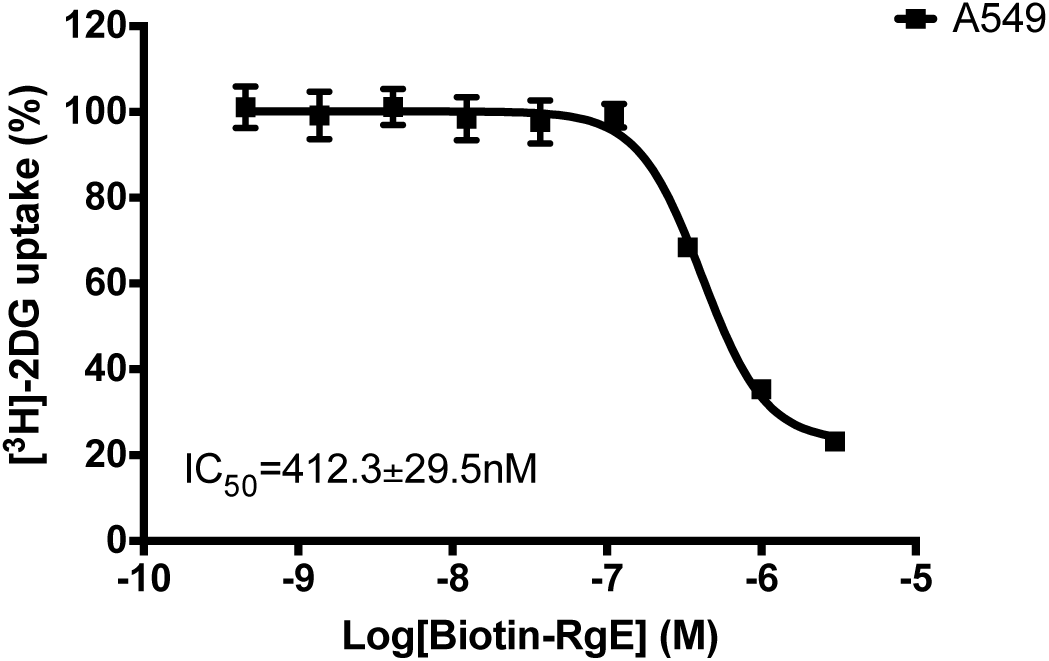
Inhibition of [^3^H]-2DG glucose uptake in A549 cells by Biotin-RgE. Error bars represent s.d.; data are mean ± s.d.; *n* = 3 independent experiments.

**Figure S5.**
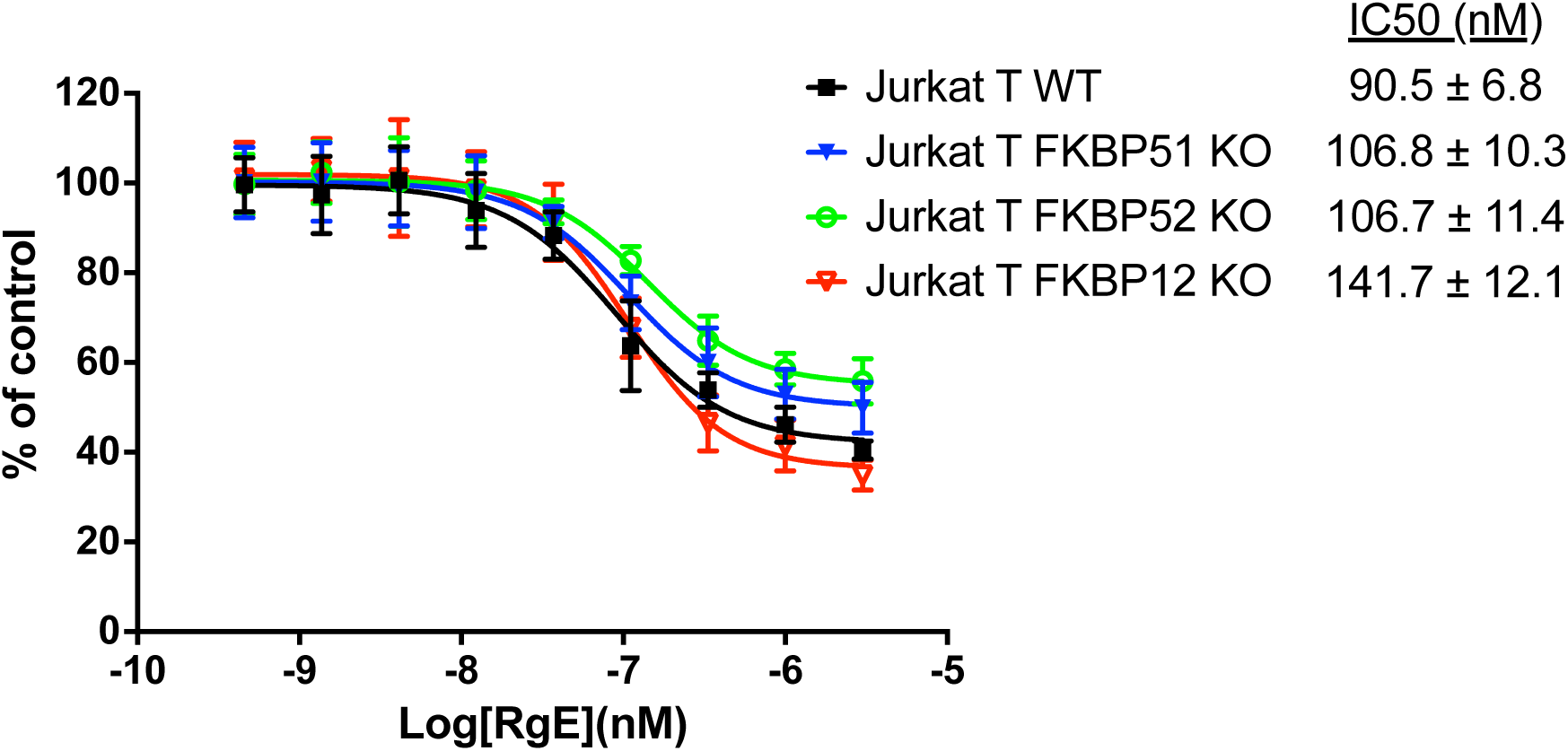
Inhibition of cell proliferation in Jurkat T wild type (WT), FKBP12 knockout (KO), FKBP51 knockout (KO), and FKBP52 knockout (KO) cells by RgE. Error bars represent s.d.; data are mean ± s.d.; *n* = 3 independent experiments.

**Figure S6.**
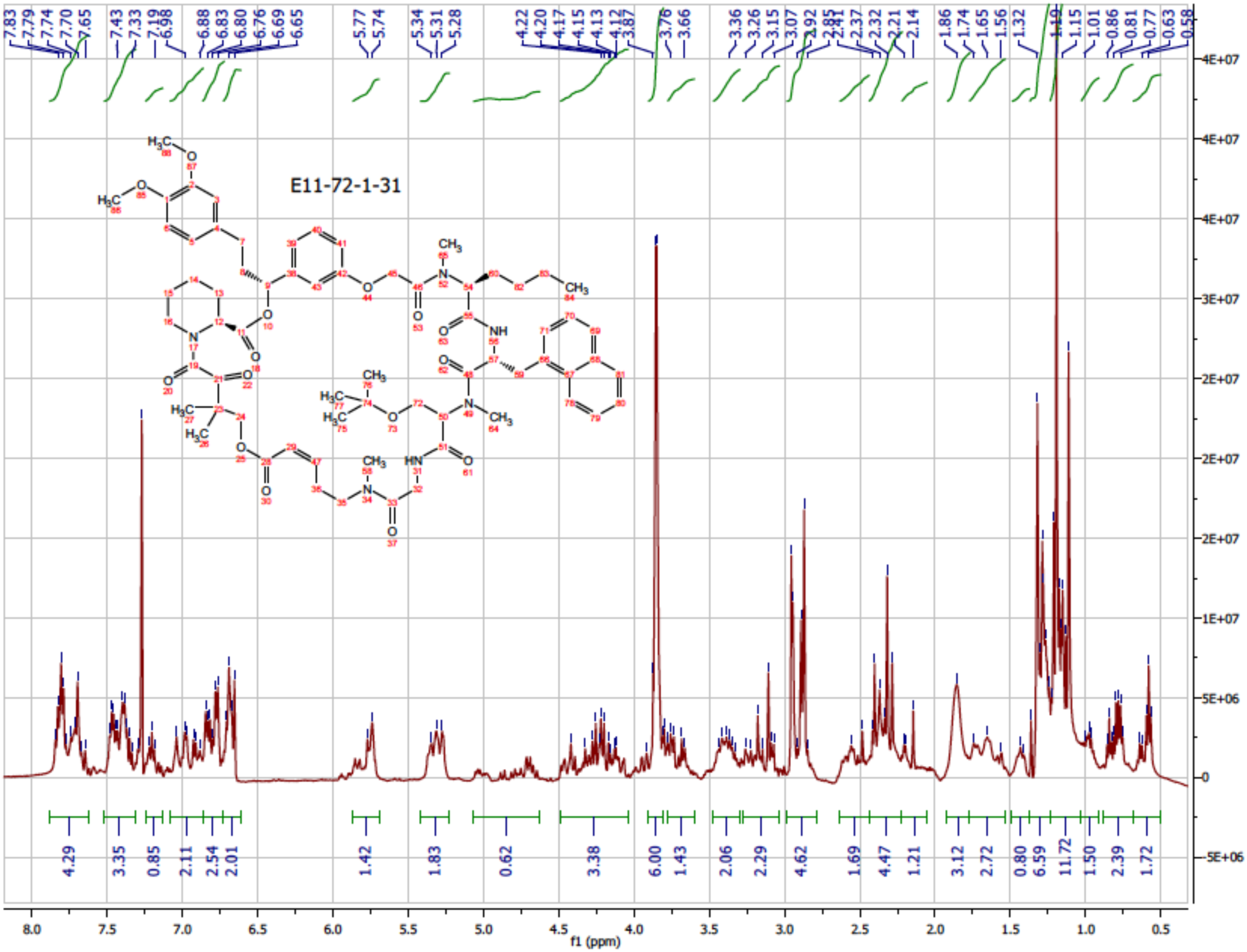
^1^HNMR of RgE (**8**).

**Figure S7.**
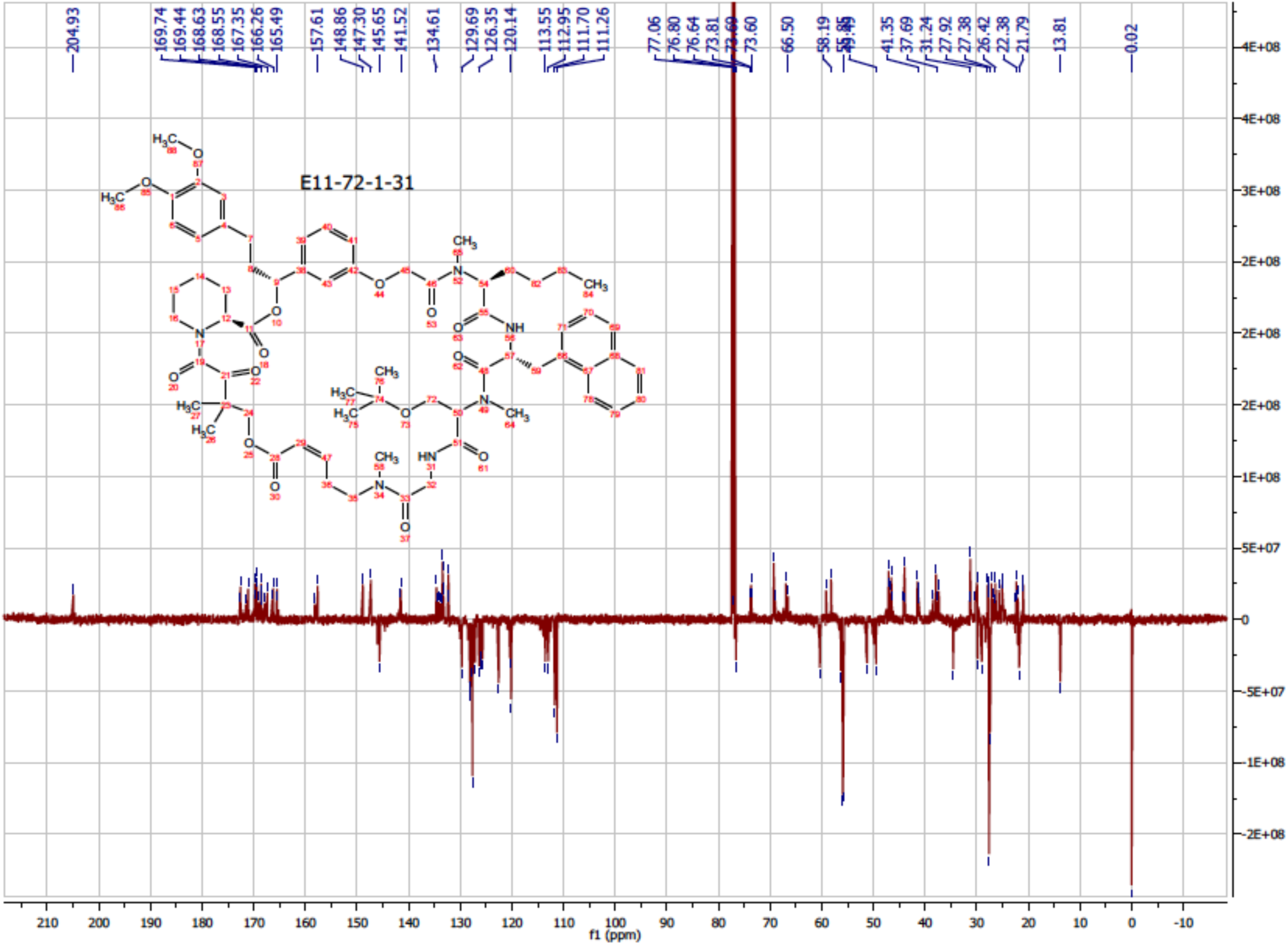
^13^C-NMR of RgE (**8**).

**Figure S8.**
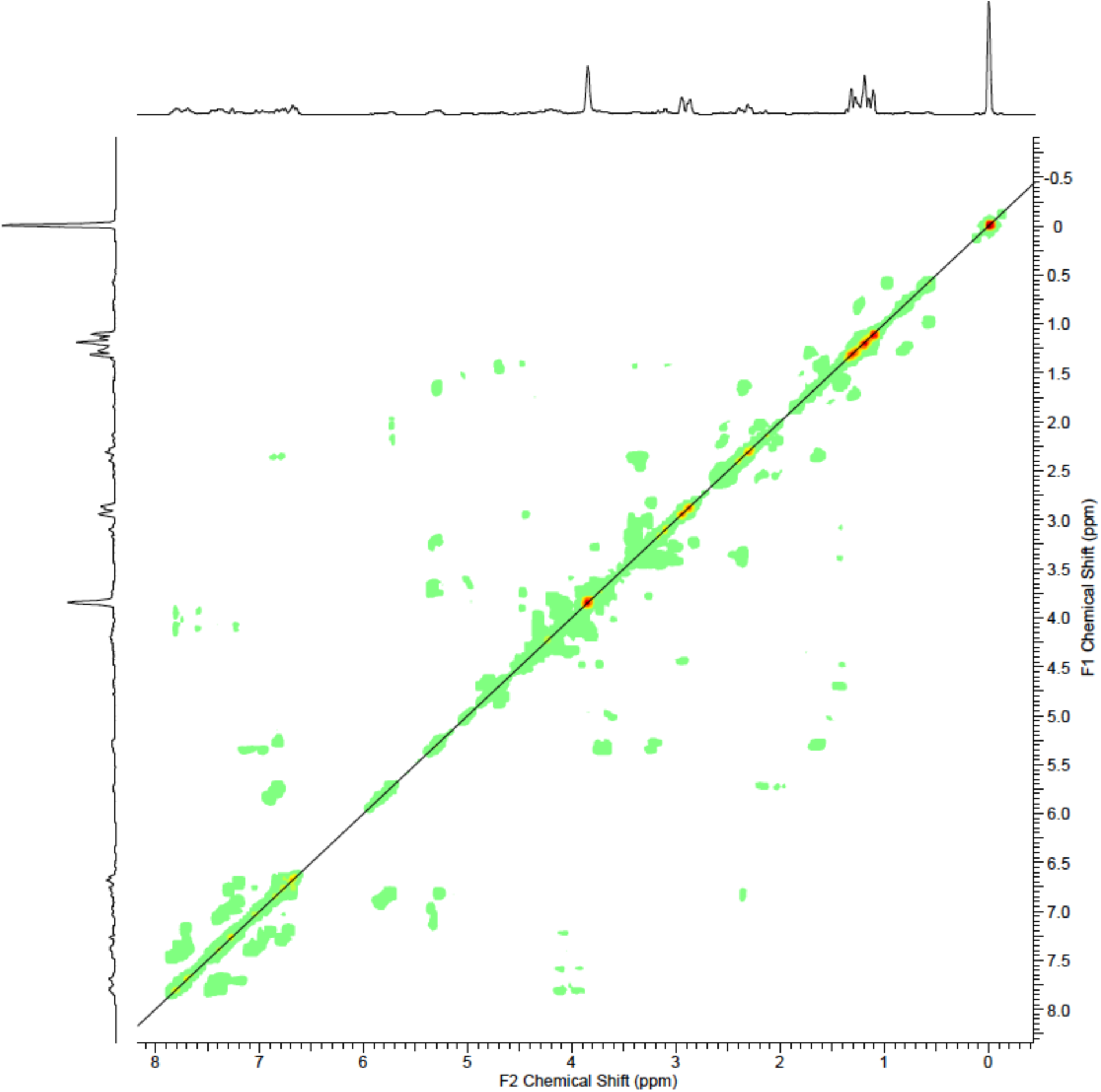
2D COSY NMR of RgE (**8**).

**Figure S9.**
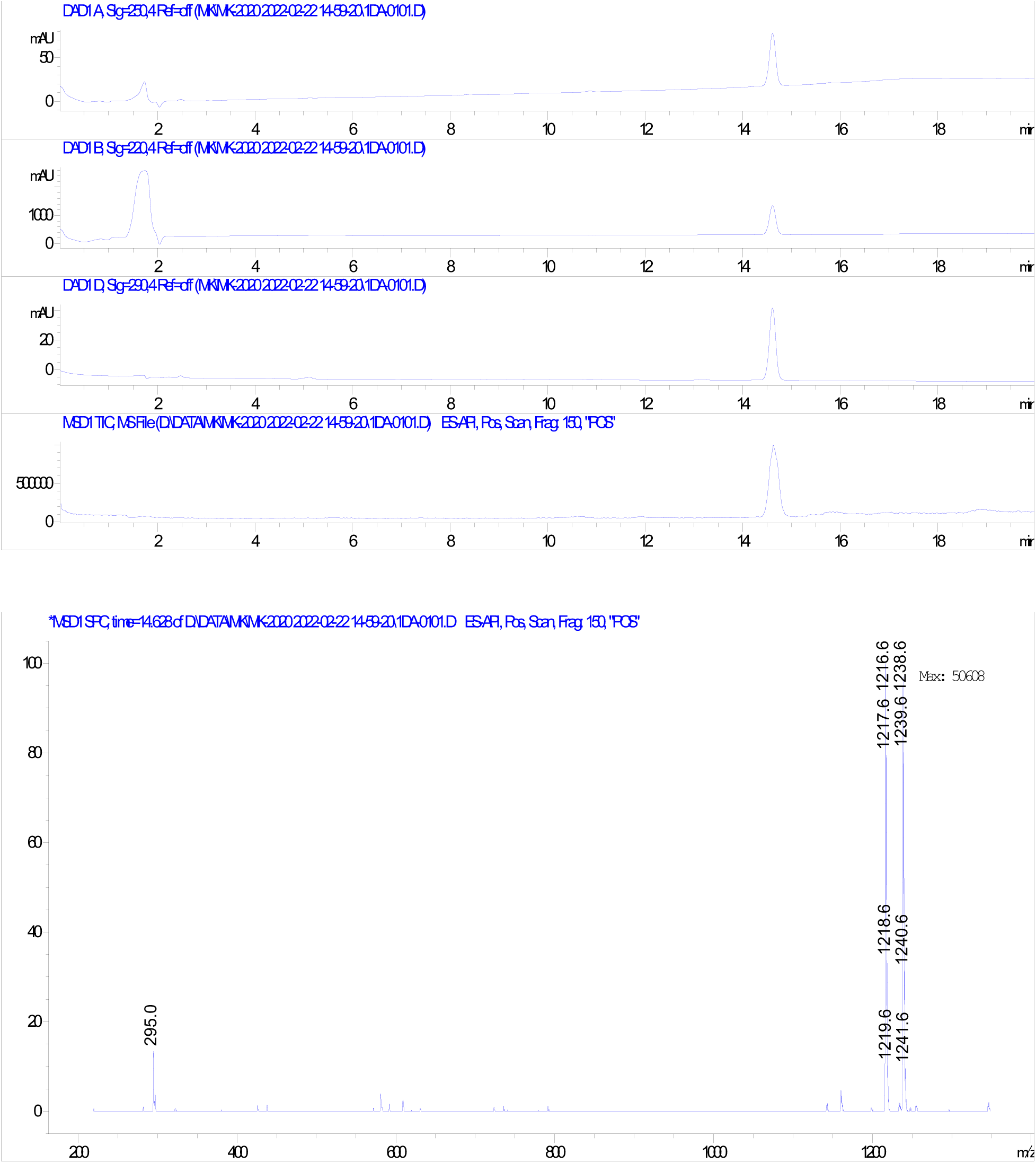
HPLC and ESI-MS of RgE (**8**).

**Figure S10.**
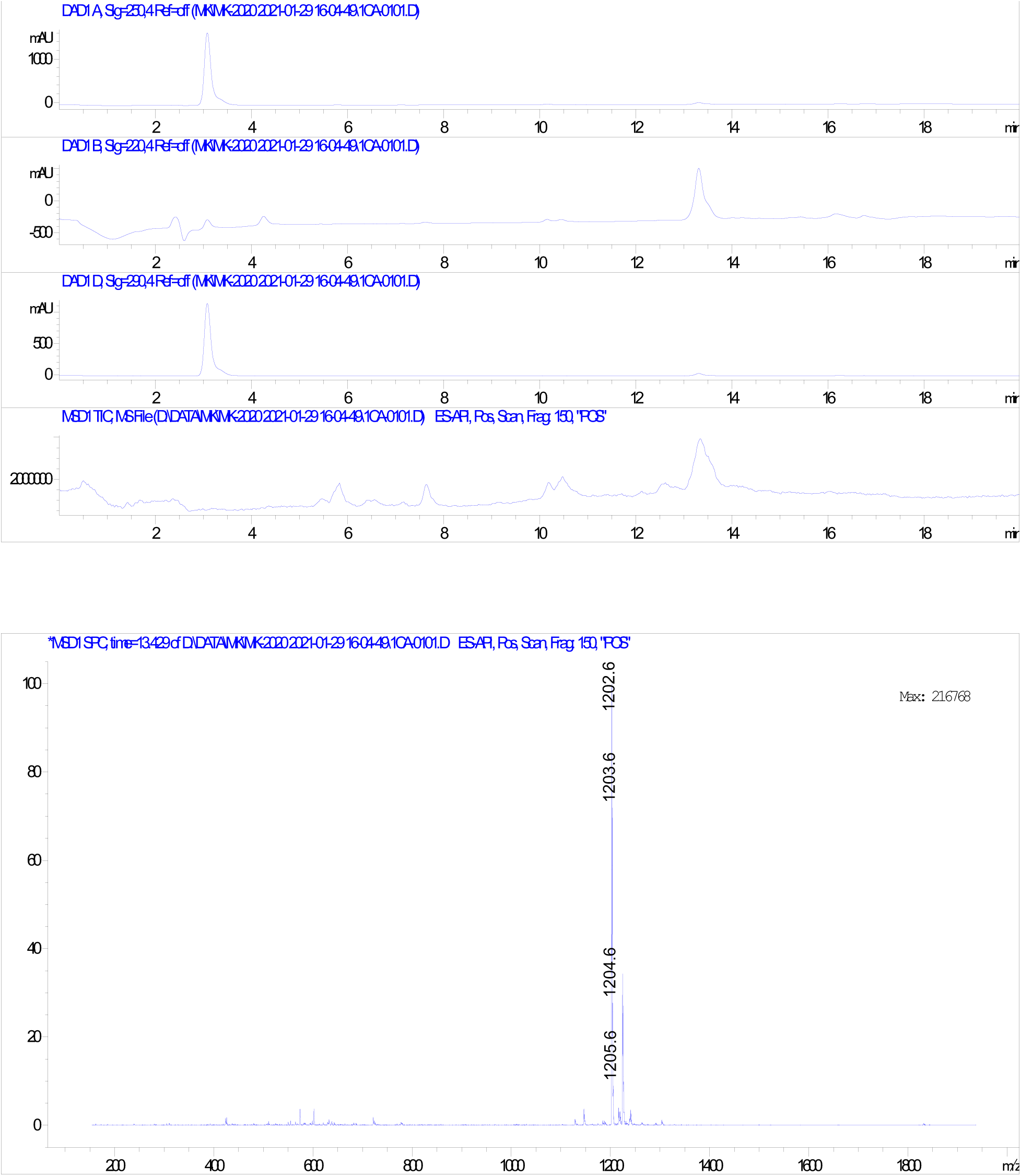
HPLC and ESI-MS of RgE-OH (**6**).

**Figure S11.**
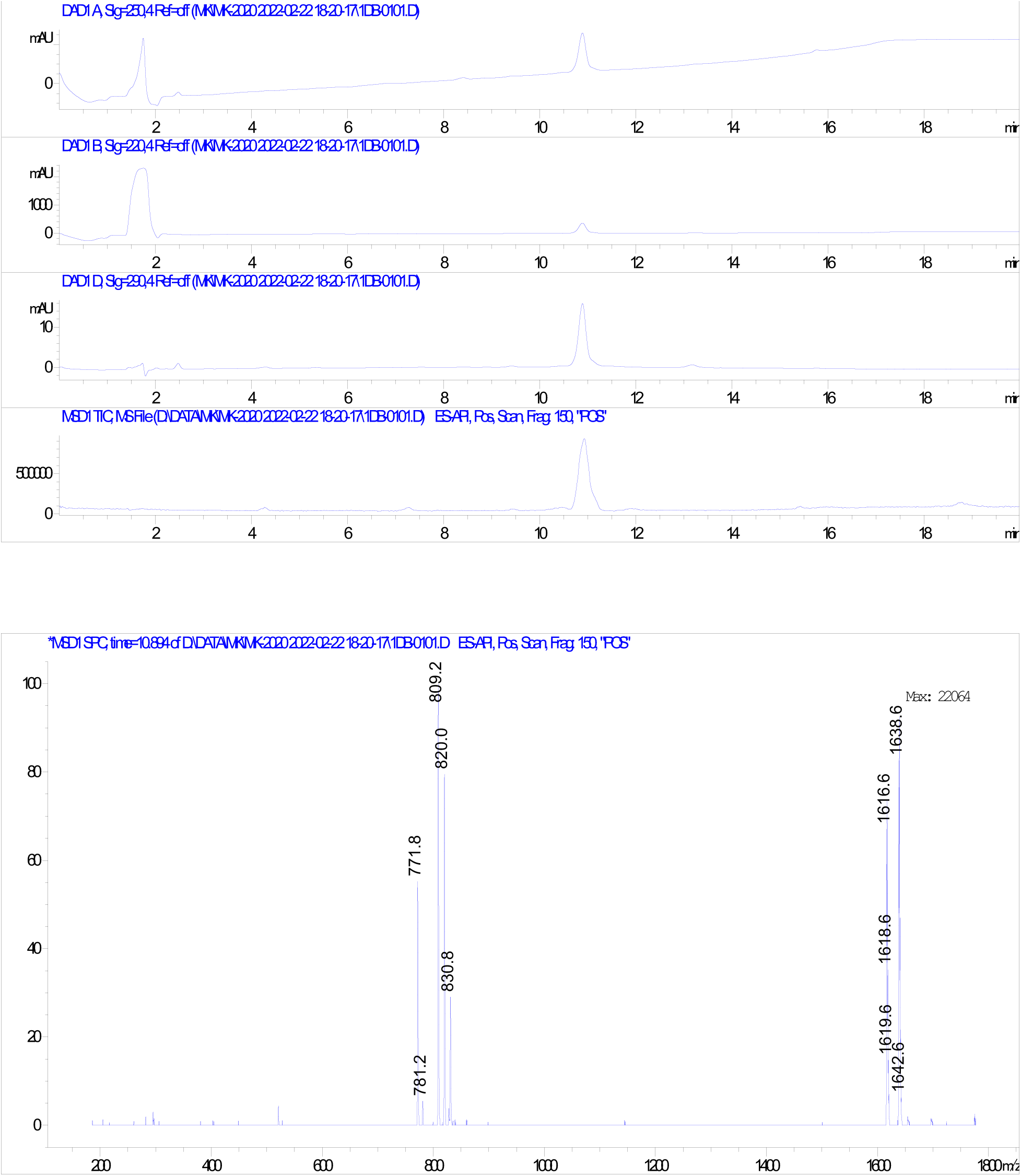
HPLC and ESI-MS of biotin-RgE (**9**).

**Table S1.** SAR study of E11-72-1 (**7**). Inhibition of 2-deoxy-D-[^3^H] glucose transport in A549 cells by E11-72-1 and its analogues. Superscript M indicates *N*-Me amino acid and D indicates D amino acids used in the sequences. Error bars represent s.d.; data are mean ± s.d.; *n* = 3 independent experiments.

| Rapafucins | AA sequences | IC<sub>50</sub> (nM) | Rapafucins | AA sequences | IC<sub>50</sub> (nM) |
| --- | --- | --- | --- | --- | --- |
| E11-72-1 | Gly-<sup>M</sup>Ser(tBu)-Nal-<sup>M</sup>Ala | 46.1±4.5 | E11-72-1-19 | Gly-<sup>M</sup>Leu-Nal-<sup>M</sup>Ala | >200 |
| E11-72-1-2 | <sup>M</sup>Gly-<sup>M</sup>Ser(tBu)-Nal-<sup>M</sup>Ala | >200 | E11-72-1-20 | Gly-<sup>M</sup>Phe-Nal-<sup>M</sup>Ala | >200 |
| E11-72-1-3 | Gly-<sup>M</sup>Ser-Nal-<sup>M</sup>Ala | >200 | E11-72-1-21 | Gly-<sup>M</sup>Tyr(tBu)-Nal-<sup>M</sup>Ala | >200 |
| E11-72-1-4 | Gly-<sup>M</sup>HoSerMe-Nal-<sup>M</sup>Ala | >200 | E11-72-1-22 | Gly-<sup>M</sup>Ser(tBu)-Tyr(tBu)-<sup>M</sup>Ala | 126.6±11.5 |
| E11-72-1-5 | Gly-<sup>M</sup>Ser(tBu)-<sup>M</sup>Phe-<sup>M</sup>Ala | >200 | E11-72-1-23 | Gly-<sup>M</sup>Ser(tBu)-PhN-<sup>M</sup>Ala | >200 |
| E11-72-1-6 | Gly-<sup>M</sup>Ser(tBu)-Phe-<sup>M</sup>Ala | >200 | E11-72-1-24 | Gly-<sup>M</sup>Ser(tBu)-Nal-<sup>M</sup>Gly | >200 |
| E11-72-1-7 | Gly-<sup>M</sup>Ser(tBu)-PhI-<sup>M</sup>Ala | 82.5±8.2 | E11-72-1-25 | Gly-<sup>M</sup>Ser(tBu)-Nal-<sup>M</sup>DAla | >200 |
| E11-72-1-8 | Gly-<sup>M</sup>Ser(tBu)-PheCl-<sup>M</sup>Ala | >200 | E11-72-1-26 | Gly-<sup>M</sup>Ser(tBu)-Nal-Pro | >200 |
| E11-72-1-9 | Gly-<sup>M</sup>Ser(tBu)-HoPhe-<sup>M</sup>Ala | >200 | E11-72-1-27 | Gly-<sup>M</sup>Ser(tBu)-Nal-<sup>M</sup>Nva | 29.2±2.5 |
| E11-72-1-10 | Gly-<sup>M</sup>Ser(tBu)-Fur-<sup>M</sup>Ala | >200 | E11-72-1-28 | Gly-<sup>M</sup>Ser(tBu)-Nal-<sup>M</sup>Phe | >200 |
| E11-72-1-12 | Gly-<sup>M</sup>Ser(tBu)-TyrOMe-<sup>M</sup>Ala | 106.1±9.8 | E11-72-1-29 | Gly-<sup>M</sup>Ser(tBu)-Nal-<sup>M</sup>Leu | >200 |
| E11-72-1-13 | Gly-<sup>M</sup>Ser(tBu)-biPhe-<sup>M</sup>Ala | >200 | E11-72-1-30 | Gly-<sup>M</sup>Ser(tBu)-Nal-<sup>M</sup>Ile | >200 |
| E11-72-1-14 | Gly-<sup>M</sup>Ser(tBu)-PhCF<sub>3</sub>-<sup>M</sup>Ala | 66.3±5.7 | E11-72-1-31 (RgE) | Gly-<sup>M</sup>Ser(tBu)-Nal-<sup>M</sup>Nle | 25.7±1.3 |
| E11-72-1-15 | Gly-<sup>M</sup>Ser(tBu)-PhpMe-<sup>M</sup>Ala | 113.5±10.6 | E11-72-1-31-2 | Gly-<sup>M</sup>Ser(tBu)-PhI-<sup>M</sup>Nle | >200 |
| E11-72-1-16 | Gly-<sup>M</sup>Ser(tBu)-Nal-Ala | >200 | E11-72-1-31-3 | Gly-<sup>M</sup>Ser(tBu)-PhCF<sub>3</sub>-<sup>M</sup>Nle | >200 |
| E11-72-1-17 | <sup>D</sup>Pro-<sup>M</sup>Ser(tBu)-Nal-<sup>M</sup>Ala | >200 | E11-72-1-31-4 | Gly-<sup>M</sup>Ser(tBu)-Tyr(tBu)-<sup>M</sup>Nle | 58.9±4.8 |
| E11-72-1-18 | Pro-<sup>M</sup>Ser(tBu)-Nal-<sup>M</sup>Ala | >200 |  |  |  |

**Table S2.** Inhibition constants of RgE (**8**) for the peptidyl prolyl cis-trans isomerase activity of different isoforms of FKBP. Error bars represent s.d.; data are mean ± s.d.; *n* = 3 independent experiments.

| Rapafucin | FKBP12<br>$K_i$ (nM) | FKBP13<br>$K_i$ (nM) | FKBP25<br>$K_i$ (nM) | FKBP51<br>$K_i$ (nM) | FKBP52<br>$K_i$ (nM) |
| --- | --- | --- | --- | --- | --- |
| RgE | 9.8 ± 2.9 | >5000 | >5000 | 93 ± 43 | 336 ± 121 |

