## Supplementary material for "Identification of Rapaglutin E as An Isoform-specific Inhibitor of Glucose Transporter 1": Materials and Methods

### Table of Contents


Materials and Methods

1. Biology

1.1 Biological Reagents

1.2 Cell culture

1.3 Over-expression of GLUT1, GLUT3, and GLUT4 in HEK293T

1.4 Affinity Pulldown with Biotinylated RgA

1.5 Western Blot Analysis

1.6 2-deoxy-D-[$^3$H] glucose ([$^3$H]-2DG) Uptake Assay

1.7 Alamar Blue Cell Viability Assay

1.8 Proteolytic foot-printing/Limited proteolysis

2. Chemistry

2.1 General Experimental for Synthesis

2.1.1 Materials and Methods

2.1.2 Synthetic Reagents

2.1.3 General Procedures: Solid-phase Peptide Synthesis (SPPS), Microwave-assisted RCM Reaction, and Macrocycle Purification protocol

2.1.4 FKBD synthesis and preparation of cis-C6 linker conjugated resin

2.2 Synthesis schemes of cis-C6 linker conjugated resin, FKBD, E11-72-1-31 (RgE) and its biotinylated probe (biotin-RgE)

2.2.1 Synthesis of cis-C6 linker conjugated resin, Pipecolic acid-acryl ester (3), and FKBD (5)

2.2.2 Scheme 1: Synthesis of OH-FKBD (4) precursor for biotin-RgE (9)

2.23 Scheme 2: Synthesis of RgE-OH (6), E11-72-1 (7), E11-72-1-31 or RgE (8) and biotin-RgE (9)

#### 2.3 Experimental Procedures and Characterization

##### 2.3.1 Experimental Procedures and Characterization of Compounds for Scheme 1

##### 2.3.2 Experimental Procedures and Characterization of Compounds for Scheme 2

#### Materials and Methods

##### 1. Biology

**1.1 Biological Reagents.** A549, HEK293T, Jurkat T E6.1, PANC10.05, and HeLa cells were sourced from ATCC and were not further authenticated. DLD1 and its *GLUT1* knockout cells were provided by Dr. Bert Vogelstein at Johns Hopkins University School of Medicine[1]. GLUT1, GLUT3, and GLUT4 overexpressing HEK 293T cells were generated and authenticated according to the method described previously[2]. Human red blood cells were obtained from ZenBio (Cat#: SER-PRBC). Jurkat T FKBP knock out cell lines were generated according to the method described previously[3]. Roswell Park Memorial Institute (RPMI) 1640, low-glucose Dulbecco's modified Eagle's medium (DMEM), and high-glucose DMEM media were purchased from Fisher Scientific (Cat#: 11875119, Cat#: 11885092, Cat#: 11995073). 2-deoxy-D-[$^3$H]-glucose was purchased from Revvity (Cat#: NET549001MC). Streptavidin agarose beads were purchased from ThermoFisher Scientific (Cat#: 20359). Antibody anti-GLUT1 was purchased from Cell Signaling Technology (Cat#: 12939).

**1.2 Cell culture.** All cells were grown at 37°C with 5% CO2 in a humidified environment. A549, HeLa, HEK293T, and DLD1 cells were cultured in low-glucose DMEM supplemented with 10% (v/v) FBS and 1% PennStrep. Jurkat T E6.1 and PANC10.05 cells were cultured in RPMI 1640 medium supplemented with 10% (v/v) FBS and 1% PennStrep. The cultures were checked periodically and found to be free of mycoplasma contamination.

**1.3 Affinity Pulldown with Biotinylated RgE and RgA.** The affinity pulldown protocol was conducted as described previously[2].

**1.4 Western Blot Analysis.** For western blot analysis, cells were harvested and lysed. Cell lysates were subjected to SDS-PAGE and then transferred to a nitrocellulose membrane. Membranes were first blocked in 5% (wt/vol) BSA in Tris-buffered saline plus 0.05% Tween 20 (TBST) at room temperature for 30 min and incubated with primary antibodies at 4 °C for overnight. Membranes were then washed three times with TBST and incubated with secondary antibodies at room temperature for 1 h. Membranes were washed with TBST three times again and incubated with ECL substrate for 2 min at room temperature. Pictures were captured using a GeneSys Image Station.

**1.5 2-deoxy-D-[$^3$H] glucose ([$^3$H]-2DG) Uptake Assay.** The inhibitory activity of compounds on glucose transport was analyzed by measuring the cell uptake of [$^3$H]-2DG as previously described[2]. Briefly, cells were washed twice and incubated in uptake buffer (100 mM NaCl, 2 mM KCl, 1 mM MgCl2, 1 mM CaCl2, 10 mM HEPES. pH 7.4) for 15 min at 37˚C. Cells were treated with drug and incubated for another 15 min before adding [$^3$H]-2DG. After 15 min incubation with [$^3$H]-2DG, cells were washed twice with uptake buffer, lysed with 0.2 N NaOH plus 0.2% SDS, and neutralized with 2M HCl. The cell lysate was transferred to a new tube containing 1 mL of Optiphase HiSafe 3 Scintillation Cocktails (1200.437, Revvity). Glucose uptake was quantified with a scintillation counter.

**1.6 Alamar Blue Cell Viability Assay.** The Alamar blue cell viability assay was conducted as described previously[2]. Cells were seeded into a 96-well plate (Costar) in 180 µL culture media. After an overnight recovery, drugs were added to make the total reaction volume 200 µL and incubated for 72h. After drug incubation, 20 µL of 0.2 mg/mL Resazurin (Sigma, Cat#: R7017) in PBS Buffer was added to cells and incubated at 37˚C for 6h before reading fluorescence (544nm Ex / 590nm Em) using a plate reader. GraphPad Prism (v4.03) software was used to determine IC50 values using a four-parameter logistic regression.

**1.7 Proteolytic foot-printing/Limited proteolysis.** The protocol is based on a method described previously[4]. GLUT1-Flag overexpressing HEK293T cells were seeded into a 10 cm dish at $1x10^6$ cells/condition in 5 mL. After an overnight recovery, cells were incubated with drug for 30 min at 37˚C. Next, media was aspirated from the plates and cells were gently rinsed in 5 mL PBS. Cells were scraped off the plate, transferred to a conical tube for centrifugation (1000 rpm for 5 min), and washed with 1 mL PBS 1x. Next, cells were resuspended in 100 µL PBS and incubated for 30 min at 37˚C with 0 or 2.5 µg pronase (Roche, Cat#: 10165921001). Following treatment, 1 mL PBS was added to dilute protease and resuspend cells that have clumped together. Next, cells were centrifuged 100 xg for 3 min and supernatant was removed. Cells were lysed in 100 µL lysis buffer (25 mM HEPES, 0.3 M NaCl, 1.5 mM $MgCl_2$, 2 mM EDTA, 2 mM EGTA, 1 mM DTT, 1% Triton X-100, and 10% glycerol buffer supplemented with protease inhibitor cocktail; DTT and protease inhibitor cocktail added fresh). After 10 min incubation on ice, samples were centrifuged at 16,000 xg for 15 min at 4˚C. The supernatants were removed and transferred to a fresh microcentrifuge tube. The protein lysate concentration was measured via the Bradford (Bio-Rad, Cat#: 5000001) protein assay. Protein lysate was mixed with sample buffer (50 mM Tris pH 6.8, 50 mM DTT, 2.5 mM EDTA, 10% glycerol, bromophenol blue; DTT added fresh) and not boiled, as high temperature induces GLUT protein aggregation. The samples were separated on 12% SDS-PAGE, transferred to a nitrocellulose membrane, and probed with anti-GLUT1 (Cell Signaling Technology, Cat#: 12939 or Abcam, Cat#: ab652) in TBST at 1:1000.

#### 2. Chemistry

##### 2.1 General Experimental for Synthesis

###### 2.1.1 Materials and Methods

Starting materials, reagents, and solvents were obtained from commercial sources with purity >95% unless otherwise noted and used as received. THF, DCM, and DMF were anhydrous grade. High performance liquid chromatographic analyses were performed with Agilent LC-MS system (Agilent 1260 series, mass detector 6120 quadrupole) with UV detection at 214, 254 nm 290 nm. Orbital shaking for solid-phase reactions was performed on a Mettler-Toledo Bohdan MiniBlock system for 96 tubes (30-200mg resin in SiliCycle tubes) or a VWR Mini Shaker (0.2-2g resin in a plastic syringe with a fritted disc). Reagents were added with an adjustable Rainin 8-channel pipette for the MiniBlock system. Microwave reactions were performed with a Biotage Initiator Plus or Multiwave Pro with silicon carbide 24-well blocks from Anton Parr.

Compound purification at 0.05-50 g scale was performed with Teledyne Isco CombiFlash Rf 200 or Biotage Isolera One system followed by a Heidolph rotary evaporator. Purification at 1-50 mg scale was performed with Agilent HPLC system. Purification of intermediates and final products was carried out on normal phase using an ISCO CombiFlash system and prepacked $SiO_2$ cartridges eluted with optimized gradients of either heptane-ethyl acetate mixture or dichloromethane-methanol as described. Columns were RediSep® Prep C18Aq Column 5 μm (30 mm x 100 mm). All target compounds had purity of >95% as established by analytical HPLC, unless otherwise noted. All purified rapafucins in the 45,000-compound library are purified in a high-throughput manner by SPE cartridges (Biotage, 460-0200-C, ISOLUTE, SI 2g/6mL) on vacuum manifold (Sigma-Aldrich, Visiprep™ SPE Vacuum Manifold, Disposable Liner, 12-port) followed by overnight drying with a custom-designed box (50 cm x 50 cm x 15 cm) that allows air flowing rapidly inside to remove the solvent. The high-throughput weighing of the compounds in the library was done by a Mettler-Toledo analytical balance that linked (Sartorious Entris line with RS232 port) to a computer with custom-coded electronic spreadsheet.

##### 2.1.2 Synthetic Reagents

Piperidine, N,N-diisopropylethylamine (DIPEA) were purchased from Alfa Aesar. Anhydrous pyridine was purchased from Across. Solid support resin with 2-chlorotrityl chloride (Cat#: 03498) was purchased from Chem-Impex. HATU was purchased from Chem-Impex. Fmoc protected amino acid building blocks were purchased from Chem-Impex, Novabiochem or GL Biochem. Dichloromethane ($CH_2Cl_2$), methanol (MeOH), hexanes, ethyl acetate (EtOAc), 1,2-dichloroethane (DCE, anhydrous), N,N'-dimethylformamide (DMF, anhydrous), $CCl_4$, methylamine (33%, ethanol), and all the other chemical reagents were purchased from Sigma-Aldrich. Hoveyda-Grubbs catalyst 2$^{nd}$ generation, Zhan 1B, Pd/C,$Pd(PPh_3)_4$ and all the other chemical reagents were purchased from Strem Chemicals. Iodoacetyl-PEG2-biotin was purchased from Biopharm.

##### 2.1.3 General Procedures: Solid-phase Peptide Synthesis (SPPS), Microwave-assisted RCM Reaction, and Macrocycle Purification protocol

*General Procedure A: Solid-phase Peptide Synthesis (SPPS)*

All orbital shaking was done at 500-600 rpm. Briefly, to prepare the resin, dry resin beads were shaking with 8x (in volume) $CH_2Cl_2$ for 20 min before the solvent was drained and washed with 1x DMF. For coupling reactions, Fmoc-protected amino acid building blocks (3.0 eq.) and HATU (3.0 eq.) were added in order into the vessel of the prepared resin. The resin and reagent mixture were quickly mixed before DIPEA (6 eq.) was added. The reaction usually took 2-3 h to completion judged by Kaiser or chloranil test. Reaction with Fmoc-valine or -isoleucine to be coupled to N-methyl amino acid on resin, the reaction mixture was drained after shaking for 3 h, briefly washed with DMF, and fresh reaction mixture was added for a second time to ensure high percentage of conversion. For FKBD conjugation, FKBD (1.6 eq.), HATU (1.6 eq.) and DIPEA (3 eq.) were used in place. Deprotection of the Fmoc group was achieved by shaking with 8x piperidine/DMF (1/4, v/v) for 20-30 min. Thorough washing was performed in between the SPPS steps by rinsing the beads with $CH_2Cl_2$ (4x) for 5 times and DMF (1x) for once.

*General Procedure B: Microwave-assisted RCM*

A total of 50 mg beads was places in a microwave reactor, followed by the addition of 1,2-dichloroethane (DCE) (5 mL/100mg) and Hoveyda-Grubbs II (30 mol%). The reactor was then sealed, and the reaction was programmed with stirring at 130 °C for 30 min under the radiation of microwave. Upon completion, the resulting brown suspension was filtered, and the filtrate was concentrated for silica gel purification ($CH_2Cl_2$: EtOAc: MeOH/ 2:2:1) or HPLC analysis.

*General Procedure C: Macrocycle Purification Protocol*

For purification using flash chromatography, 4 g ISCO zip columns were equilibrated in 100% $CH_2Cl_2$. The samples taken from the microwave reactors were dissolved in minimal DCM and injected using a filtered syringe onto the columns. The flash purification subjected the samples to a linear gradient from 0% methanol in $CH_2Cl_2$ and EtOAc to 20% methanol in $CH_2Cl_2$ and EtOAc over 60 column volumes (CV). After 60 CVs, the gradient was set back to 0% methanol so the columns could be equilibrated for the next sample. Each column was used 5 times before discarding. LC-MS Analytical Protocol for Macrocycles Analytical reversed-phase high-performance liquid chromatography (HPLC) was performed on a C-18 reverse phase HPLC column (5μm Luna, 0.46 cm × 25 cm). Separations were achieved using linear gradients of buffer B in A (A = 0.1% formic acid in $H_2O$; B = 0.1% formic acid in $CH_3CN$) at a flow rate of 1 ml/min.

#### 2.2 Synthesis schemes of cis-C6 linker conjugated resin, FKBD, E11-72-1-31 (RgE) and biotin-RgE conjugate

##### 2.2.1 Synthesis of cis-C6 linker conjugated resin, (*S*)-1-(4-(acryloyloxy)-3,3-dimethyl-2-oxobutanoyl)piperidine-2-carboxylic acid (3), and FKBD (5).

See method previously described[5].

##### 2.2.2 Scheme 1: Synthesis of OH-FKBD (4) precursor for biotin-RgE

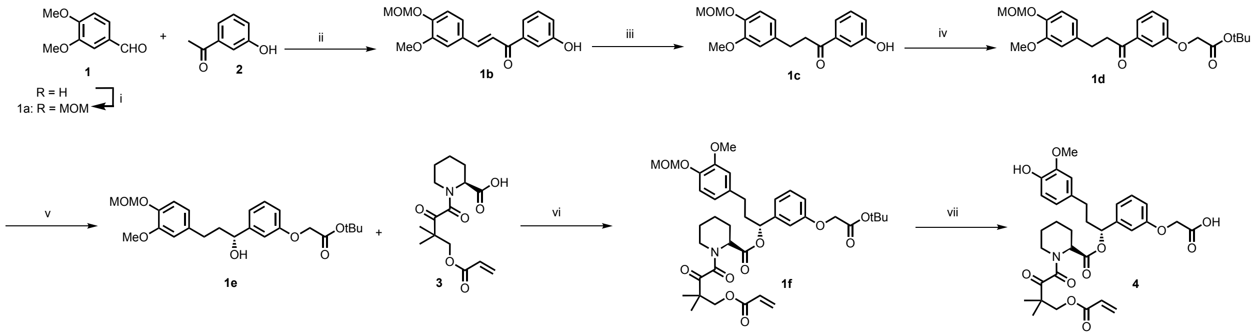


Scheme 1. *Reagents conditions:* (i) aq. KOH, $H_2O$/EtOH (1/20), RT, 12 h; (ii) Pd/C (10%), $H_2$, MeOH, RT, 1-3 h; (iii) tert-butyl bromoacetate, $K_2CO_3$, DMF, RT, 8 h; (iv) (+)-DIPCl, THF, -20 °C 16 h.

##### 2.2.3 Scheme 2: Synthesis of RgE-OH (6), E11-72-1 (7) RgE (8), and biotin-RgE (9)

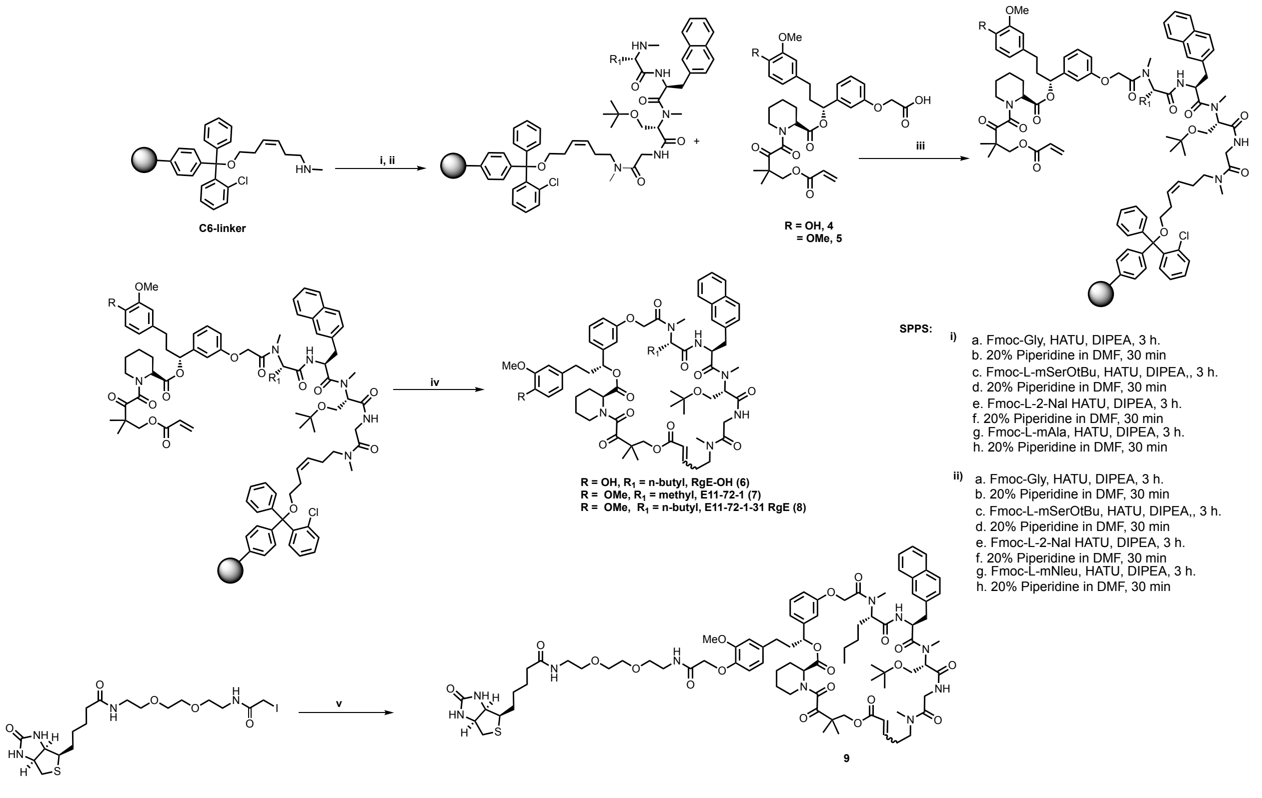


Scheme **2**. *Reaction conditions*: (i) & (ii) SPPS (iii) HATU, DIEA, DMF, 4 h; (iv) Zhan 1B, DCE, 120 °C, 1 h; (v) (**6**), $K_2CO_3$, DMF, RT, 24 h.

#### 2.3 Experimental Procedures and Characterization of Compounds

##### 2.3.1 Experimental Procedures and Characterization of Compounds for Scheme 1

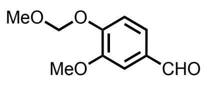

**3-methoxy-4-(methoxymethoxy)benzaldehyde (1a):** To a solution of vanillin (10 g, 1 eq, 65.78 mmol) in dry $CH_2Cl_2$ (100 mL) were added MOMCl (7.94 g, 7.49 mL, 1.5 eq, 98.6 mmol) and DIPEA (12.73 g, 17 mL, 1.5 eq, 98.68 mmol) at 0 °C under Ar atmosphere. After stirring for 4 h at room temperature, the reaction was quenched by adding saturated aqueous $NaHCO_3$ at 0 °C. The aqueous layer was extracted three times with $CH_2Cl_2$. The combined organic layers were dried over $MgSO_4$, filtered, and evaporated under reduced pressure. The crude product was purified by flash chromatography (petroleum/EtOAc = 6/1) to give (**1a**) as a white solid (9 g, 90%). $^1$H-NMR (500 MHz, $CDCl_3$): δ 9.87 (s, 1H), 7.43 (d, *J* = 8.0 Hz, 2H), 7.27 (d, *J* = 9.5 Hz, 1H), 5.32 (s, 2H), 3.95 (s, 3H), 3.52 (s, 3H). ESI-MS: m/z 197.21[M+H]$^+$.

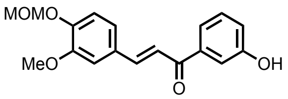


**(*E*)-1-(3-hydroxyphenyl)-3-(3-methoxy-4-(methoxymethoxy)phenyl)prop-2-en-1-one (1b):** MOM protected vanillin (**1a**) (8.0 g, 1 eq, 40.81 mmol) and 3-hydroxy acetophenone (**2**) (5.82 g, 1.05 eq, 42.8 mmol) mixture in EtOH (100 mL), was cooled to 0 ºC and treated with a cold of aq. KOH (4.57 g, 2 eq, 81.6 mmol in 5 mL water). The resulting solution was allowed to warm to room temperature (rt) and stirred for 12 h. The resulting slurry of yellow precipitate reaction mixture was concentrated then neutralized with 1N HCl (pH = 1) and then dissolved in EtOAc (100 mL) and washed with water (2×200 mL). Washed with brine (2×200 mL), dried over $Na_2SO_4$, filtered, and concentrated to afford chalcone product (**1b**) is pure enough for the next step. (11.5 g, 83.2%). TLC (EtOAc/hexane, 1/3) Rf=0.4; $^1$H-NMR (500 MHz, $CDCl_3$): δ 7.75 (d, *J* = 15.6 Hz, 1H), 7.61 (s, 1H), 7.56 (d, *J* = 7.6 Hz, 1H), 7.46 (s, 1H), 7.39-7.34 (m, 2H), 7.28-7.26 (m, 1H), 7.10 (d, *J* = 9.6 Hz, 1H), 6.92 (d, *J* = 8.4 Hz, 1H), 6.30 (s, 1H), 5.29 (s, 2H), 3.93 (s, 3H), 3.55 (s, 3H). ESI-MS: m/z 337.33[M+Na]$^+$.

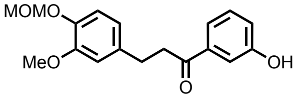


**1-(3-hydroxyphenyl)-3-(3-methoxy-4-(methoxymethoxy)phenyl)propan-1-one (1c):**
To a solution of α,β-unsaturated ketone (**1b**) (9.0 g, 28.66 mmol) in MeOH (60 mL), Pd/C (10%, 1 g) was added. The reaction vessel was flushed with hydrogen gas repetitively by using a balloon of hydrogen and high vacuum. The reaction mixture was stirred at RT for 1-3 h before filtered through a pad of celite. The filtrate was concentrated and subject to column chromatography (80 g silica gel) and eluted with EtOAc/hexane (1:3). Purified (**1c**) was collected as a pale yellow solid (7.9 g, 87.7%). TLC (EtOAc/hexane, 1/1): Rf=0.4; $^1$H NMR (500 MHz, $CDCl_3$): δ 7.47 (d, *J* = 7.7 Hz, 1H), 7.41 (s, 1H), 7.30 (t, *J* = 7.9 Hz, 1H), 7.04 (d, *J* = 7.3 Hz, 2H), 6.84-6.81 (m, 2H), 6.27 (s, 1H), 5.21 (s, 2H), 3.84 (s, 3H), 3.53 (s, 3H), 3.22 (t, *J* = 7.6 Hz, 3H), 2.97 (t, *J* = 7.6 Hz, 3H). ESI-MS: m/z 339.39[M+Na]$^+$.

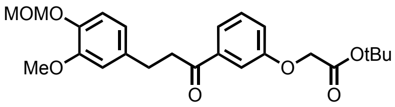


***tert*-butyl 2-(3-(3-(3-methoxy-4-(methoxymethoxy)phenyl)propanoyl)phenoxy)acetate (1d):**
A solution of the corresponding phenol (**1c**) (7 g, 1 eq, 22.15 mmol) in dry DMF (100 ml) under an atmosphere of nitrogen was treated with $K_2CO_3$ (4.58 g, 1.5 eq, 33.22 mmol) and tert-butyl bromoacetate (5.183 g, 3.956 mL, 1.2 eq, 26.56 mmol). The mixture was stirred at room temperature for 8 h until TLC indicated complete conversion. The precipitate of the title compound was collected by filtration, washed with water (3 x 200 ml) and dissolved in EtOAc,

concentrated crude product. Purified through flash chromatography and eluted with EtOAc/hexane (1:3). Purified (**1d**) was collected as a colorless liquid (8.3 g, 68.14%). $^1$H-NMR (500 MHz, $CDCl_3$): δ 7.56 (d, *J* = 7.6 Hz, 1H), 7.46 (s, 1H), 7.36 (t, *J* = 7.9 Hz, 1H), 7.13 (d, *J* = 8.3 Hz, 1H), 7.04 (s, 1H), 6.86-6.81 (m, 2H), 5.22 (s, 2H), 4.56 (s, 2H), 3.86 (s, 3H), 3.52 (s, 3H), 3.24 (t, *J* = 7.7 Hz, 2H), 2.98 (t, *J* = 7.7 Hz, 2H), 1.49 (s, 9H); ESIMS: m/z 431.47 [M+H]+.

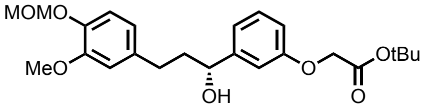


***tert*-butyl(*R*)-2-(3-(1-hydroxy-3-(3-methoxy-4-(methoxymethoxy)phenyl)propyl)phenoxy)acetate (1e):** A solution of ketone (**1d**) (6 g, 1eq, 13.9 mmol) was dried under high-vacuum for 1 h and the reaction vessel was flushed with argon and dissolved in Dry THF in 75 mL) was added and the solution was cooled down to -20 °C. (+)-DIPCl (1.6M in hexane, 3.5 eq., 15.9 mL) was added via a syringe and the resulting yellow solution was allowed to stand in a -20 °C freezer for 16 h before being warmed up to RT. Stirring at RT for 30 min, and the reaction mixture was quenched with 2,2'-(ethylenedioxy)diethylamine (1.0 eq to DIPCl) by forming an insoluble complex. After stirring at RT for another 30 min, the suspension was filtered through a pad of celite and concentrated. Column chromatography (80-200 mesh) with EtOAc/hexane (1/1) afforded product (**1e**) (5.1 g, 85%) as a colorless oil. $^1$H-NMR (500 MHz, $CDCl_3$): δ 7.28-7.25 (m, 1H), 7.05 (d, *J* = 8.2 Hz, 1H), 6.96 (d, *J* = 7.4 Hz, 1H), 6.93 (s, 1H), 6.83-6.79 (m, 1H), 6.72-6.67 (m, 2H), 5.19 (s, 2H), 4.68-4.64 (m, 1H), 4.52 (s, 2H), 3.86 (s, 3H), 3.51 (s, 3H), 2.72-2.56 (m, 2H), 2.12-1.94 (m, 2H), 1.48 (s, 9H); ESI-MS: m/z 433.55 [M+H]+ .

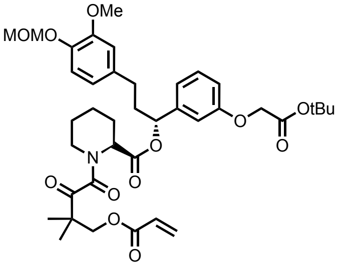


**(*R*)-1-(3-(2-(*tert*-butoxy)-2-oxoethoxy)phenyl)-3-(3-methoxy-4-(methoxymethoxy)phenyl)propyl(*R*)-1-(4-(acryloyloxy)-3,3-dimethyl-oxobutanoyl)piperidine-2-carboxylate (1f):** To solution of asymmetric alcohol (**1e**) (3 g, 1 eq, 7.45 mmol) and pipecolic acid (**3**)[5] (2.16 g, 1 eq, 6.95 mmol) were dissolved in a mixture of anhydrous DCM (30 mL) and $Et_3N$ (2.10 g, 2.89 mL, 3 eq, 20.79 mmol) and DMAP (85 mg, 0.1 eq, 0.694 mmol) were added in order and stirred for few minutes at 0 °C. To this mixture, benzoyl chloride (0.976 g, 0.806 mL, 1 eq, 6.94 mmol) was added and the resulting suspension was stirred at room temperature for 4-6 h until TLC indicated complete conversion. The reaction mixture was diluted with water and extracted three times with dichloromethane. The combined organic layers were washed three times with water and once with brine, dried over magnesium sulfate and concentrated to an oil. The crude title compound was subject to column chromatography (80 g mesh) with EtOAc/hexane (1:1). (3.8 g, 73.6%) was collected as a white foam (**1f**). $^1$H-NMR (500 MHz, $CDCl_3$): δ 7.28-7.25 (m, 2H), 7.03 (d, *J* = 8.0 Hz, 1H), 6.95 (d, *J* = 7.7 Hz, 1H), 6.92-6.87 (m, 1H), 6.82 (d, *J* = 8.3 Hz, 1H), 6.70-6.63 (m, 2H), 6.37 (d, *J* = 18.6 Hz, 1H), 6.04 (dd, *J* = 17.3, 10.4 Hz, 1H), 5.81 (d, *J* = 9.2 Hz, 1H), 5.78-5.75 (m, 1H), 5.28 (s, 1H), 5.18 (s, 2H), 4.53-4.51 (m, 3H), 4.35 (d, *J* = 11.0 Hz, 1H), 4.27-4.24 (m, 1H), 3.85 (s, 3H), 3.50 (s, 3H), 3.18-3.12 (m, 1H), 2.60-2.46 (m, 2H), 2.36 (d, *J* = 13.6 Hz, 1H), 2.26-2.19 (m, 2H), 2.08-2.01 (m, 2H), 1.77-1.65 (m, 2H), 1.61 (d, *J* = 11.3 Hz, 1H), 1.47 (s, 9H), 1.34 (s, 6H). $^{13}$C-NMR (126 MHz, $CDCl_3$) δ 205.1, 169.6, 168.0, 166.6, 165.7, 158.3, 149.8, 144.9, 141.4, 135.4, 131.5, 129.9, 128.1, 120.5, 120.0, 116.8, 114.4, 113.5, 112.3, 95.8, 82.5, 77.5, 77.2, 77.0, 76.9, 69.4, 65.9, 56.3, 56.0, 53.6, 51.5, 46.8, 44.1, 38.0, 31.4, 28.2, 26.6, 25.2, 22.4, 21.7, 21.3. ESI-MS calculated for $C_{39}H_{52}NO_{12}$ 726.84 $[M+H]^+$ observed: 726.38.

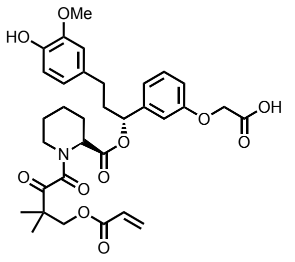

**2-(3-((*R*)-1-(((*R*)-1-(4-(acryloyloxy)-3,3-dimethyl-2-oxobutanoyl)piperidine-2-carbonyl)oxy)-3-(4-hydroxy-3-methoxyphenyl)propyl)phenoxy)acetic acid (4):**
Compound (**1f**) (2 g, 1 eq, 2.5 mmol) was dissolved in 30 mL of dichloromethane in a round-bottom flask under Ar protection. Then 20% TFA (6 mL) was added through a syringe in 3 portions during 6 h while stirring at room temperature. The reaction was monitored through TLC. When full conversion was achieved, solvents and TFA were removed under vacuum. Product was purified by column chromatography (80-200 mesh) with 5-10% MeOH/DCM. FKBD (**4**) (1.6 g, 80%) was collected as a light-yellow foam. $^1$H-NMR (500 MHz, $CDCl_3$): δ 7.29-7.25 (m, 1H), 6.94 (d, *J* = 7.7 Hz, 1H), 6.87 (d, *J* = 6.3 Hz, 1H), 6.81 (d, *J* = 7.9 Hz, 1H), 6.66-6.61 (m, 2H), 6.36 (d, *J* = 17.3 Hz, 1H), 6.04 (dd, *J* = 17.3, 10.5 Hz, 1H), 5.81 (d, *J* = 10.5 Hz, 1H), 5.77-5.72 (m, 1H), 5.28 (d, *J* = 5.4 Hz, 1H), 4.67 (d, *J* = 7.3 Hz, 2H), 4.33 (d, *J* = 11.2 Hz, 1H), 4.26 (d, *J* = 11.0 Hz, 1H), 3.86 (s, 3H), 3.46 (d, *J* = 13.0 Hz, 1H), 3.22-3.17 (m, 1H), 2.62-2.49 (m, 1H), 2.38 (d, *J* = 14.2 Hz, 1H), 2.27-2.20 (m, 1H), 1.80-1.76 (m, 1H), 1.74-1.70 (m, 1H), 1.62 (d, *J* = 13.0 Hz, 1H), 1.48 (dt, *J* = 13.0, 3.7 Hz, 1H), 1.32 (d, *J* = 3.2 Hz, 6H). $^{13}$C-NMR (126 MHz, $CDCl_3$) δ 204.9, 171.8, 171.5, 169.4, 166.8, 165.9, 158.0, 146.7, 144.1, 141.9, 132.8, 131.6, 130.1, 128.0, 121.1, 120.2, 115.2, 114.5, 111.8, 111.1, 69.5, 65.2, 60.7, 56.1, 51.8, 46.9, 44.2, 38.3, 31.6, 26.7, 25.1, 22.3, 21.7, 21.3, 21.3, 14.4. ESIMS calculated for $C_{33}H_{39}NO_{11}Na$ 648.67 $[M+Na]^+$ observed: 648.33.

##### 2.3.2 Experimental Procedures and Characterization for Scheme 2

For E11-72-1 (**7**), Fmoc protected Glycine, N-methyl SerOtBu, 2-Napthylalanine, N-methyl alanine were coupled in order on to cis-C6 linker conjugated beads (General Procedure A) before microwave-assisted RCM reaction (General Procedure B). For RgE (**8**) and RgE-OH (**6**), Fmoc protected Glycine, N-methyl SerOtBu, 2-Napthylalanine, N-methyl norleucine, and FKBDs were coupled in order on to cis-C6 linker conjugated beads (General Procedure A) before microwave-assisted RCM reaction (General Procedure B). For E11-72-1 (**7**), RgE (**8**), and RgE-OH (**6**), silica gel purification method was followed in General Procedure C to yield 30-45% at ~50 mg scale.

**(12S,4R,10S,13S,16S)-16-(tert-butoxymethyl)-10-butyl-4-(3,4-dimethoxyphenethyl)-9,15,21,29,29-pentamethyl-13-(naphthalen-2-ylmethyl)-3,6,27-trioxa-9,12,15,18,21-pentaaza-1(2,1)-piperidina-5(1,3)-benzenacyclohentriacontaphan-24-ene-2,8,11,14,17,20,26,30,31-nonaone (7):** For E11-72-1 (**7**) Fmoc protected Glycine, N-methyl SerOtBu, 2-Napthylalanine, N-methyl alanine, and FKBD (**5**) were coupled in order on to cis-C6 linker conjugated beads (General Procedure A) before microwave-assisted RCM reaction (General Procedure B). For RgE **7** silica gel purification method was followed in General Procedure C to yield 30-45% at ~50 mg scale. $^1$H-NMR (500 MHz, $CDCl_3$): δ 7.83-7.79 (m, 2H), 7.74-7.65 (m, 2H), 7.48-7.42 (m, 1H), 7.40-7.31 (m, 2H), 7.19 (t, *J* = 7.9 Hz, 1H), 7.10-6.96 (m, 1H), 6.91-6.83 (m, 1H), 6.76 (d, *J* = 8.4 Hz, 2H), 6.70-6.66 (m, 2H), 6.64 (s, 1H), 5.85-5.78 (m, 1H), 5.76-5.71 (m, 1H), 5.34-5.28 (m, 2H), 5.05-4.96 (m, 1H), 4.48-4.38 (m, 1H), 4.32-4.12 (m, 2H), 3.85 (s, 6H), 3.77-3.66 (m, 2H), 3.51-3.22 (m, 3H), 3.17-3.07 (m, 2H), 2.94 (s, 3H), 2.87 (d, *J* = 8.6 Hz, 3H), 2.62-2.46 (m, 2H), 2.40-2.28 (m, 4H), 2.16 (s, 2H), 1.83-1.63 (m, 3H), 1.54-1.40 (m, 1H), 1.29 (d, *J* = 18.2 Hz, 9H), 1.20 (d, *J* = 6.2 Hz, 8H), 1.14-1.09 (m, 3H), 0.84-0.75 (m, 3H), 0.62-0.55 (m, 2H) ppm. $^{13}$C-NMR (126 MHz, $CDCl_3$) δ 204.9, 169.7, 169.4, 168.6, 168.5, 167.3, 166.2, 165.4, 157,6, 148.8, 147.3, 145.6, 141.5, 134.6, 129.6, 126.3, 120.1, 113.5, 112.9,111.7,111.2, 73.6,66.5, 58.1, 55.8, 49.4, 41.3, 37.6, 31.2, 27.9, 27.3, 26.4, 22.3, 21.7, 13.8 ppm. ESI-MS calculated for $C_{64}H_{82}N_6O_{15}$ = 1174.58 $[M+H]^+$ observed: 1175.6.

**(12S,4R,10S,13S,16S)-16-(tert-butoxymethyl)-4-(3,4-dimethoxyphenethyl)-9,10,15,21,29,29-hexamethyl-13-(naphthalen-2-ylmethyl)-3,6,27-trioxa-9,12,15,18,21-pentaaza-1(2,1)-piperidina-5(1,3)-benzenacyclohentriacontaphan-24-ene-2,8,11,14,17,20,26,30,31-nonaone (8):** For E11-72-1-31 or RgE (**8**) Fmoc protected Glycine, N-methyl SerOtBu, 2-Napthylalanine, N-methyl norleucine and FKBD (**5**) were coupled in order on to cis-C6 linker conjugated beads (General Procedure A) before microwave-assisted RCM reaction (General Procedure B). For RgE (**8**) silica gel purification method was followed in General Procedure C to yield 30-45% at ~50 mg scale. ESI-MS calculated for $C_{67}H_{88}N_6O_{15}$ = 1216.83 $[M+H]^+$ observed: 1218.6.

**(12S,4R,10S,13S,16S)-16-(tert-butoxymethyl)-10-butyl-4-(4-hydroxy-3-methoxyphenethyl)-9,15,21,29,29-pentamethyl-13-(naphthalen-2-ylmethyl)-3,6,27-trioxa-9,12,15,18,21-pentaaza-1(2,1)-piperidina-5(1,3)-benzenacyclohentriacontaphan-24-ene-2,8,11,14,17,20,26,30,31-nonaone (6):** For RgE-OH (**6**) Fmoc protected Glycine, N-methyl SerOtBu, 2-Napthylalanine, N-methyl norleucine, and FKBD (**4**) were coupled in order on to cis-C6 linker conjugated beads (General Procedure A) before microwave-assisted RCM reaction (General Procedure B). For RgE-OH (**6**) silica gel purification method was followed in General Procedure C to yield 30-45% at ~50 mg scale. ESI-MS calculated for $C_{66}H_{86}N_6O_{15}$ = 1202.62 $[M+H]^+$ observed: 1203.6.

**Biotinylated-E11-72-31 or biotin-RgE** (**9**)**:** A solution of the RgE-OH (**6**) (20 mg, 1 eq, 0.016mmol) in dry DMF (200 µl) under an atmosphere of nitrogen was treated with $K_2CO_3$ (23 mg, 10 eq, 0.166 mmol) and 13.5 mg iodoacetyl-PEG2-biotin (13.5 mg, 1.5 eq, 0.025mmol). The mixture was stirred at room temperature for 24 h until complete conversion. The precipitate of the title compound was centrifuge and collected crude product. Purified through Prep HPLC and collected as 2 mg, by 10% yield of pure biotin-RgE (**9**). ESI-MS calculated for $C_{84}H_{117}N_{10}O_{20}S$ = 1616.81 $[M+H]^+$observed: 1618.6.
